# A Gata4–Loxl2 axis controls ECM remodeling to enable scar-free spinal cord regeneration in zebrafish

**DOI:** 10.64898/2026.01.21.700220

**Authors:** Nicolas P. Noel, Vânia F. Lima Fernandes, Andrew Kjar, Paige E. Pfotenhauer, Mawusi P. Sefogbe, Eman A. Akam-Baxter, Ethan S. Lippmann, Valentina Cigliola

## Abstract

Spinal cord injury in mammals causes permanent loss of sensory and motor function due to scar formation obstructing axon regrowth. In contrast, zebrafish fully regenerate their spinal cord without lasting fibrosis. Here, we identify the transcription factor Gata4 as a key positive regulator of regeneration. After injury, *gata4* expression is predominantly upregulated in ependymo-radial glial cells and in glial cells extending processes toward sites of tissue bridging. Gata4 preserves glial identity and orchestrates regenerative extracellular matrix (ECM) remodeling. Gata4 loss biases glial cells toward a fibroblast-like state with upregulated Loxl2b, a collagen cross-linking enzyme, and can also increase Loxl2b in nearby non-glial cells. Excess Loxl2b activity promotes pathological crosslinking and formation of a rigid, fibrotic matrix impairing glial and axonal bridging across the lesion, greatly reducing motor recovery. Pharmacological inhibition of Loxl2b in *gata4* mutants substantially rescues ECM organization, tissue bridging, and functional recovery. Injury-induced *gata4* expression depends, at least in part, on Hif1α signaling and activation of a short injury-responsive enhancer. We identify the Gata4–Loxl2b axis as a central regulator of and potential therapeutic target for scar-free spinal cord repair.

## INTRODUCTION

Spinal cord injury (SCI) causes profound neuronal loss, axonal disruption, and synapse elimination, leading to permanent sensory, motor, and autonomic deficits in mammals. Beyond the initial mechanical insult, secondary injury responses including immune cell infiltration, activation of resident glia and recruitment of mesenchymal cell types, drive the formation of a multicellular injury scar^1,2^. This scar consists of a fibrotic lesion core surrounded by a glial border, and it fundamentally shapes the post-injury environment encountered by regenerating axons^1,2^. The fibrotic lesion core forms due to excessive extracellular matrix (ECM) deposition by invading non-neural cells, particularly platelet-derived perivascular fibroblasts. This dense matrix alters biochemical and mechanical properties of the tissue, thereby restricting axonal regrowth^3–5^. Reactive astrocytes and microglia at the lesion border stabilize the injury, support restoration of the blood–spinal cord barrier, and form a glial boundary that segregates spared tissue from the lesion^1,2^. Astrocytes also deposit inhibitory ECM components, including chondroitin-sulfate proteoglycans and tenascin-C, at this border, further shaping the inhibitory post-injury environment^6–8^. Reducing ECM deposition from fibroblasts or astrocytes, or selectively removing inhibitory ECM components, increases lesion permissiveness and facilitates axonal regeneration^1,5,9,10^.

In contrast to mammals, zebrafish possess a remarkable capacity to regenerate the spinal cord and fully recover motor function throughout life^11–14^. Following SCI in zebrafish, fibroblasts deposit sparse ECM and selectively limit inhibitory components, such as small leucine-rich proteoglycans, while the injury environment contains a transient, regeneration-permissive ECM^15,16^. *pdgfrb⁺* myoseptal and perivascular fibroblasts are a major source of injury-associated ECM^17^. Some of these cells biphasically regulate inflammation by interacting with innate immune cells, first promoting and then resolving the acute inflammatory phase, establishing a regeneration-permissive lesion environment^18^. In addition, ependymo-radial glial cells (ERGs), progenitor cells lining the central canal, proliferate after injury with a subset contributing to the formation of a glial bridge across the lesion site, which has been proposed to provide structural support for axon regrowth^13,19^. Together, sparse inhibitory ECM, transient regenerative ECM, glial bridging, and tightly controlled inflammation create an environment that actively supports axon regeneration. However, the transcriptional programs linking fibroblast and glial responses to ECM remodeling and a regeneration-permissive niche remain largely unknown.

GATA4, a member of the GATA family of transcription factors (TFs), has emerged as a potent regulator of tissue repair, mostly working in a facilitating manner. The vertebrate GATA family comprises six TFs sharing a conserved two zinc-finger DNA-binding domain recognizing the consensus motif (A/T)GATA(A/G) or W-GATA-R. GATA4 has received extensive attention for its role in tissue regeneration, particularly in the heart. It is induced in zebrafish cardiomyocytes after ventricular resection injury and marks a population that drives myocardial regeneration^20^. In neonatal mice, GATA4 is similarly required for heart repair, as its deletion impairs regeneration, reduces cardiomyocyte proliferation, and increases scar size^21^. Mechanistically, GATA4 controls expression of cell-cycle genes (e.g., Cyclin A2, Cyclin D2 and CDK4) and promotes pro-regenerative paracrine factors such as FGF16^21,22^. Beyond the heart, GATA4 also regulates tissue remodeling and anti-fibrotic responses in the liver. In hepatic stellate cells, GATA4 promotes fibrosis regression by repressing fibrogenic genes and activating anti-fibrotic transcriptional programs, while endothelial GATA4 prevents pathogenic remodeling and supports tissue regeneration^23–26^. Emerging evidence indicates that GATA4 is expressed in neurons and glia in the adult central nervous system (CNS) and regulates astrocyte proliferation and apoptosis in vitro^27^. Its loss in human astrocytes is associated with malignant transformation, suggesting that GATA4 may influence glial cell reactivity, proliferation, and perhaps regenerative responses in the CNS^28^. Thus, GATA4 emerges as a candidate transcriptional orchestrator of proliferation, cell-state plasticity, and repair across multiple organs, including the CNS.

Here, we identify Gata4 as a central regulator of spinal cord regeneration in adult zebrafish, acting primarily in ERGs and in glial cells extending processes toward sites of tissue bridging, and to a lesser extent in cells surrounding the lesion site. Following injury, Gata4 is strongly upregulated and controls expression of the lysyl oxidase family enzyme Loxl2b, which normally contributes to ECM structure. Loss of Gata4 function increases Loxl2b levels, leading to excessive collagen cross-linking that stiffens the matrix and creates an anti-regenerative environment, impairing axon growth and functional recovery. Injury-induced *gata4* expression depends, at least in part, on Hif1α signaling and a short upstream enhancer that harbors Hif1α binding sites. These findings establish Gata4 as a key transcriptional regulator linking injury-induced gene expression to ECM remodeling in adult zebrafish, primarily through control of ECM-modifying enzymes such as Loxl2b, which determine the extent of ECM deposition during spinal cord regeneration.

## RESULTS

### *gata4* regulatory sequences are activated during spinal cord regeneration

In a screen for regulators of zebrafish spinal cord regeneration, we identified *gata4* regulatory sequences as being strongly activated after injury. In situ hybridization revealed negligible endogenous *gata4* transcripts in the uninjured spinal cord, in contrast to a sharp increase in *gata4*-expressing cells around the central canal and a moderate increase throughout the spinal cord following transection injury **(Figure S1A)**. To visualize this response more precisely, we examined spinal cords of *Tg(−14.8kb-gata4:EGFP)* zebrafish, carrying a *gata4* upstream regulatory region extensively shown to recapitulate the endogenous *gata4* expression pattern^29^. In uninjured tissue, enhanced green fluorescent protein (EGFP) expression was largely undetectable, and sporadically limited to neurons identified by expression of the HuC/D marker **(Figures 1A and S1B)**. At 1 week post-injury (wpi), *gata4*-driven EGFP was detected near the rostral and caudal injury stumps, in cells scattered along the central canal, and in a small population of flattened cells surrounding the spinal cord, likely corresponding to myoseptal and perivascular fibroblasts **(Figures 1A and 1B**, yellow arrows and arrowheads, respectively). EGFP signal intensified and peaked at 2 wpi, with EGFP-positive cells present at sites of tissue bridging, and decreased by 4 wpi, essentially coinciding with the completion of regeneration **(Figure 1A)**. Cross-sectional analyses at 1 and 2 wpi showed *gata4*-driven EGFP expression primarily confined to ERGs surrounding the central canal and dorsal spinal parenchyma **(Figures 1B and 1C)**.

**Figure 1.**
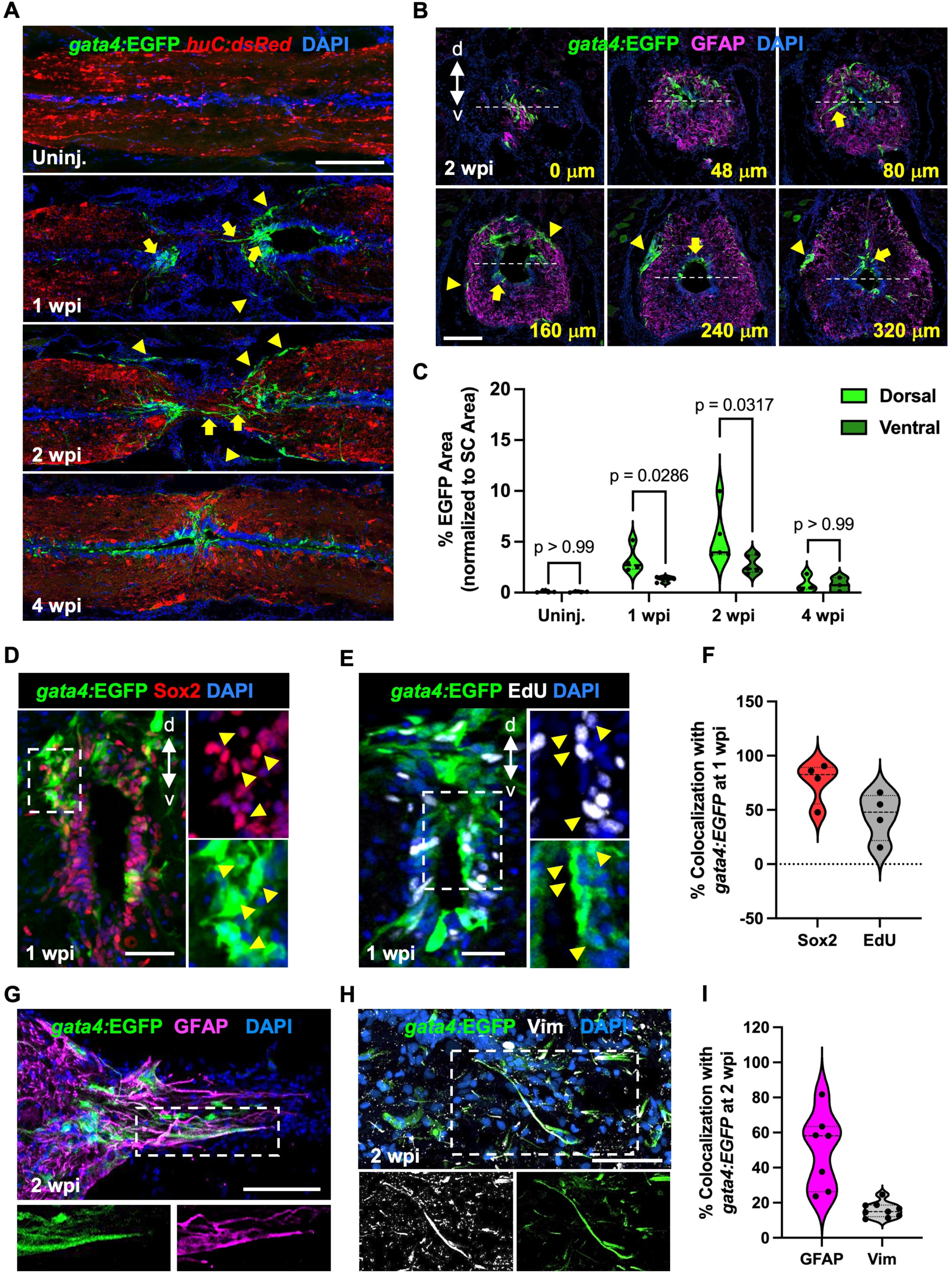
Gata4 regulatory sequences are induced during spinal cord regeneration. **(A)** Sections of spinal cord tissue indicating *Tg(−14.8kb-gata4:*EGFP) reporter expression (green) in uninjured, 1, 2 and 4 weeks post-injury (wpi) animals. *Tg(huC:*dsRED) marks neurons. Yellow arrows and arrowheads indicate EGFP^+^ cells in and around the regenerating site, respectively. **(B)** Immunofluorescence staining showing expression of *gata4*:EGFP (green) and the glial marker GFAP (magenta) in cross sections along the proximodistal axis of the regenerating spinal cord. Dashed lines delineate mid points of the central canal along the dorso-ventral axis. Yellow values indicate distance from the injury site. **(C)** Quantification of *Tg(−14.8kb-gata4:*EGFP) reporter signal in dorsal and ventral central canal and spinal parenchyma domains at 2 wpi. Dorsal and ventral regions were defined relative to the central canal midline. SC, spinal cord. **(D)** Assays for Sox2 (red) expression at 1 wpi in transverse sections of *Tg(−14.8kb-gata4:*EGFP) spinal cord. **(E)** EdU (white) incorporation assay for cell cycling at 1 wpi in transverse sections of *Tg(−14.8kb-gata4:*EGFP) spinal cord. **(F)** Percentage of *gata4*-expressing cells positive for Sox2 or EdU at 1 wpi. **(G)** Longitudinal section of *Tg(−14.8kb-gata4:*EGFP) spinal cord at 1 wpi showing *gata4* expression in glial cells visualized using GFAP (magenta) and **(H)** in vimentin-positive cells (white). **(I)** Percentage of *gata4*-expressing cells positive for GFAP or Vimentin at 2 wpi. Dashed area in **D**, **E**, **G** and **H** indicate magnified regions. Scale bars 200 μm in **A**, 100 μm in **B**, 50 μm in **D**, **E**, **G** and **H**. d dorsal, v ventral.

To define the identity of *gata4*-expressing cells during regeneration, we conducted histological and molecular analyses using antibodies and transgenic markers. At 1 wpi, 76% of Gata4⁺ cells co-expressed Sox2, a marker of ERGs **(Figures 1D and 1F)**, and 44.3% showed indicators of cell cycling based on 5-ethynyl-2′-deoxyuridine (EdU) incorporation assays **(Figures 1E and 1F)**. By 2 wpi, roughly 49.9% of Gata4^+^ cells expressed the glial marker GFAP **(Figures 1G, 1I, S1C and S1D)**, while 15.8% expressed vimentin, a mesenchymal marker previously shown to be expressed in glia acquiring a migratory phenotype^30^ **(Figures 1H and1I)**. As GFAP immunostaining primarily labels glial processes rather than cell bodies, the degree of GFAP colocalization is likely underestimated. Gata4^+^ cells co-expressing Olig2⁺, marking progenitors associated with oligodendrocyte and motor neuron lineages^12^, were observed very rarely at 2 wpi **(Figure S1E)**.

All six GATA transcription factors remain poorly characterized functionally in the CNS. We probed *gata* gene expression in a published transcriptomic dataset of GFAP^+^ glial subpopulations at 1 wpi^31^ and found that *gata3*, *gata4*, *gata5*, and *gata6* were expressed across different glial clusters after injury. Notably, *gata4* was only upregulated in clusters 1 and 2 **(Figure S1F)**, consistent with expression in a majority, but not all, glial cell processes observed by histology **(Figures 1G, S1C and S1D)**. Overall, our data identify *gata4* as a regeneration-responsive gene predominantly expressed in ERGs and glial cells extending processes at sites of tissue bridging.

### Gata4 loss-of-function impairs progenitor responses and delays spinal cord regeneration

To evaluate the requirement for Gata4 during regeneration, we used *gata4^wcm6/wcm6^* zebrafish, carrying a 6-bp TALEN-mediated insertion that introduces a premature stop codon at the start of the C-terminal GATA4 zinc-finger domain, eliminating DNA-binding capability^32^ **(Figures 2A, 2B and S2A)**. Uninjured *gata4^wcm6/wcm6^* embryos appeared phenotypically normal and survived to adulthood without overt morphological abnormalities, although *gata4* null mutants were recovered at sub-Mendelian frequencies as both embryos and adults, as previously shown^32^.

**Figure 2.**
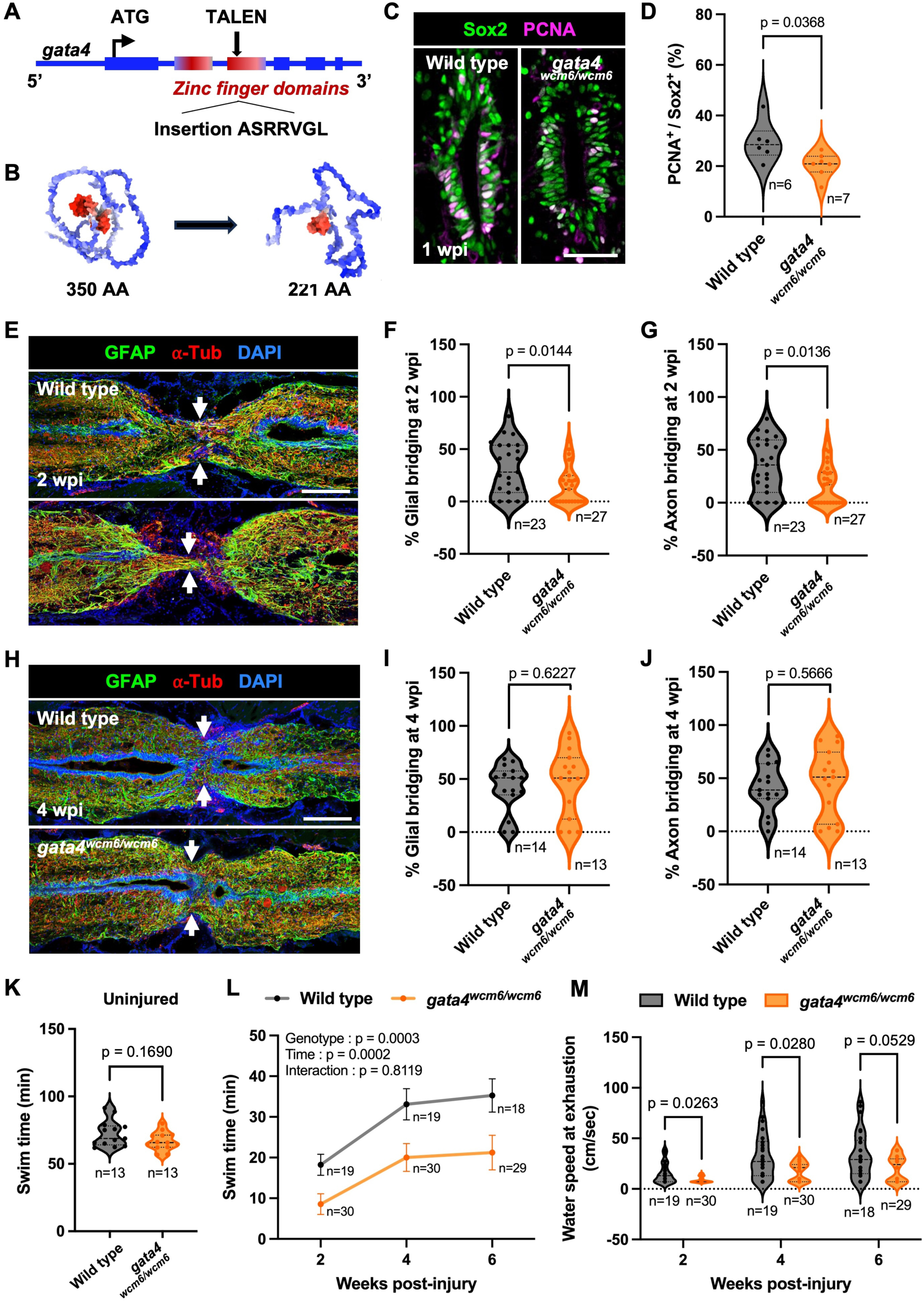
Gata4 loss-of-function delays tissue bridging and impairs motor recovery after spinal cord injury. **(A)** Schematic showing the TALEN insertion in the second *gata4* zinc finger domain used to generate the *gata4^wcm6/wcm6^* zebrafish line. **(B)** Alpha Fold representation of *gata4^wcm6/wcm6^* and wild-type Gata4 protein, highlighting predicted structural differences and protein length. AA, amino acid. **(C)** Ependymal cell proliferation assessed by PCNA staining in spinal cords of *gata4^wcm6/wcm6^* and wild-type zebrafish on transverse sections at 1 week post-injury (wpi). Quantification shown in **(D). (E)** Tissue sections stained for the glial marker GFAP (green) and the axonal marker acetylated α-Tubulin (red) in wild-type and *gata4^wcm6/wcm6^* spinal cords at 2 wpi. Arrows indicate tissue bridging. Quantification of glial bridging and axon bridging shown in **(F)** and **(G)**, respectively. **(H)** Tissue sections stained for the glial marker GFAP (green) and axonal marker acetylated α-Tubulin (red) in wild-type and *gata4^wcm6/wcm6^* spinal cord at 4 wpi. Arrows indicate sites of tissue bridging. Quantification of glial bridging and axon bridging shown in **(I)** and **(J)**, respectively. **(K)** Endurance swim testing of uninjured wild-type and *gata4^wcm6/wcm6^* zebrafish. Animals were placed in an enclosed swim tunnel with increasing water current, and the time to exhaustion was recorded. **(L)** Endurance swim testing of injured wild-type and *gata4^wcm6/wcm6^* zebrafish at 2, 4, and 6 wpi. **(M)** Time-course of water current velocity at which *gata4^wcm6/wcm6^* and wild-type zebrafish could no longer maintain forward swimming. Scale bar 50 μm in **C**, 100 μm in **E** and **H**. Welch’s *t*-test used for comparisons in **D**; Mann-Whitney used for comparisons in **F**, **G**, **I**, **J, K, M**; Two-Way ANOVA used for comparisons in **L**.

We subjected *gata4^wcm6/wcm6^* animals to SCI and histologically and functionally assessed regeneration at multiple time-points. *gata4^wcm6/wcm6^* animals showed a nearly 30% reduction in Sox2^+^ ERG proliferation at 1 wpi compared to wild-type clutch-mates, assessed by PCNA staining on cross-sections (p = 0.0368, Welch’s *t*-test) **(Figures 2C and 2D)**. We then examined the glial and axonal bridging across the lesion site on longitudinal spinal cord sections at 2 wpi, using GFAP as a glial marker, and acetylated α-tubulin as an axonal marker. The average diameter of the glial and axonal bridge was reduced by 52.1% and 51.4%, respectively, in *gata4^wcm6/wcm6^* animals compared to wild type **(Figures 2E-G and S2B)**. These defects were no longer detectable by 4 wpi, when tissue bridge measurements in *gata4^wcm6/wcm6^* animals equaled those of wild types **(Figures 2H-J)**.

To assess spinal cord function, we quantified swim capacity using a ramped-current swim challenge in which fish swam against progressively increasing water current velocity until failure. The time to failure and the corresponding current velocity were recorded as quantitative measures of swim capacity. While no significant difference in swim capacity was detected between uninjured *gata4^wcm6/wcm6^* and wild-type animals **(Figure 2K)**, *gata4^wcm6/wcm6^* fish exhibited significantly reduced swimming endurance at 2, 4 and 6 weeks following SCI **(Figures 2L and S2C)**. Although swimming speed of *gata4^wcm6/wcm6^* animals improved over time, it remained considerably lower than in wild-type animals **(Figure 2M)**. Strikingly, despite recovery of tissue bridging by 4 wpi **(Figures 2H–J)**, swim defects persisted **(Figures 2L and 2M)**, suggesting that factors beyond tissue bridging limit motor function recovery or that a finer level of functional or circuit-level analysis is required to capture subtle deficits. Overall, these results demonstrate that Gata4 function is required for effective spinal cord regeneration and full recovery of motor function.

### Loss of Gata4 function shifts glial transcriptional programs toward a fibrotic state

To further characterize the *gata4^wcm6/wcm6^* phenotype, we performed single-nucleus RNA sequencing (snRNA-seq) of wild-type and *gata4^wcm6/wcm6^* spinal cords at 2 wpi, when *gata4* expression is maximal **(Figures 3A and S3A)**. Unsupervised clustering of 4,626 nuclei identified 20 cell clusters, including major spinal cord cell types, fibroblasts, and immune cells, distinguished by specific hallmark genes **(Figures 3B and S3C)**. Cell-cluster composition was similar between genotypes, with no substantial differences in the relative proportions of cell types **(Figure S3B)**. We next used volcano plots to visualize transcriptional differences between *gata4^wcm6/wcm6^* and wild-type ERGs/Glia (hereafter referred to as glia, excluding oligodendrocytes and microglia), as these cells were most enriched in *gata4* expression post-injury **(Figure 3C)**. These comparisons revealed widespread transcriptomic changes in glia after *gata4* loss-of-function, with gene-expression profiles differing markedly between genotypes **(Table S1)**. Notably, one of the most strongly upregulated genes in *gata4^wcm6/wcm6^* glia was *lysyl oxidase 2b (loxl2b),* which encodes an extracellular enzyme involved in collagen and elastin crosslinking, and ECM remodeling **(Figure 3C)**. Gene-set enrichment analysis (GSEA) of *gata4^wcm6/wcm6^* glia revealed significant enrichment of ECM-related genes, including those involved in ECM–receptor interactions **(Figure 3D and Table S2)**. No enrichment of these genes was observed in oligodendrocyte, immune, or neural clusters, and the fibroblast cluster had insufficient cell numbers for comparison between *gata4^wcm6/wcm6^* and wild-type spinal cords **(Figures S3D-F)**. The upregulated genes in glia included multiple collagen components (*col4a1, col9a2, col1a1b, col18a1a, col6a1, col2a1b, col4a2, col4a5, col4a4, col4a6, col1a2, col2a1a, col1a1a, col6a4a, col6a2, col27a1b and col4a3*), which encode structural elements of the basal lamina and fibrillar ECM that influence tissue architecture and stiffness. Additionally, genes for laminin subunits (*lamb4, lamb2*) and integrin receptors (*itga10, itga9, itga6a, itga6b, itgb1a*), which mediate cell–ECM adhesion and signaling, were also over-represented. Several matrix-associated proteins (*frem1a, thbs4b, thbs1b, cc44b*) were likewise increased in *gata4^wcm6/wcm6^* glia compared to wild type **(Figures 3E, S4A and S4B)**. We validated expression of three ECM-related transcripts by *in situ* hybridization on spinal cord sections of *gata4^wcm6/wcm6^* and wild types. In uninjured spinal cords, *loxl2b*, *col181a1* and *col27a1b* showed basal expression in cells surrounding the central canal **(Figure S5A)**. Following injury, these transcripts were upregulated in and around the lesion site in *gata4^wcm6/wcm6^* spinal cord compared to wild type **(Figure 3F-N)**. Glial expression of *loxl2b* was further confirmed by performing *in situ* hybridization with a *loxl2b*-specific probe on spinal cord sections of *Tg(−14.8kb-gata4:EGFP)* **(Figure S5B).** The significant enrichment of the ‘ECM–receptor interaction’ category suggests that the *gata4^wcm6/wcm6^* mutation may alter cell-matrix signaling, potentially modifying the composition or organization of the extracellular matrix in a way that impairs tissue bridging. In agreement with that, increased *loxl2b* has been associated with enhanced matrix stiffness and fibrotic responses in several organs after injury^33,34^. This observation, together with the observed increase in ECM-related genes in *gata4^wcm6/wcm6^* versus wild type, suggests that *gata4* loss-of-function may shift glia and derived glia programs toward an increased ECM-remodeling state or fibrotic transcriptional profile.

**Figure 3.**
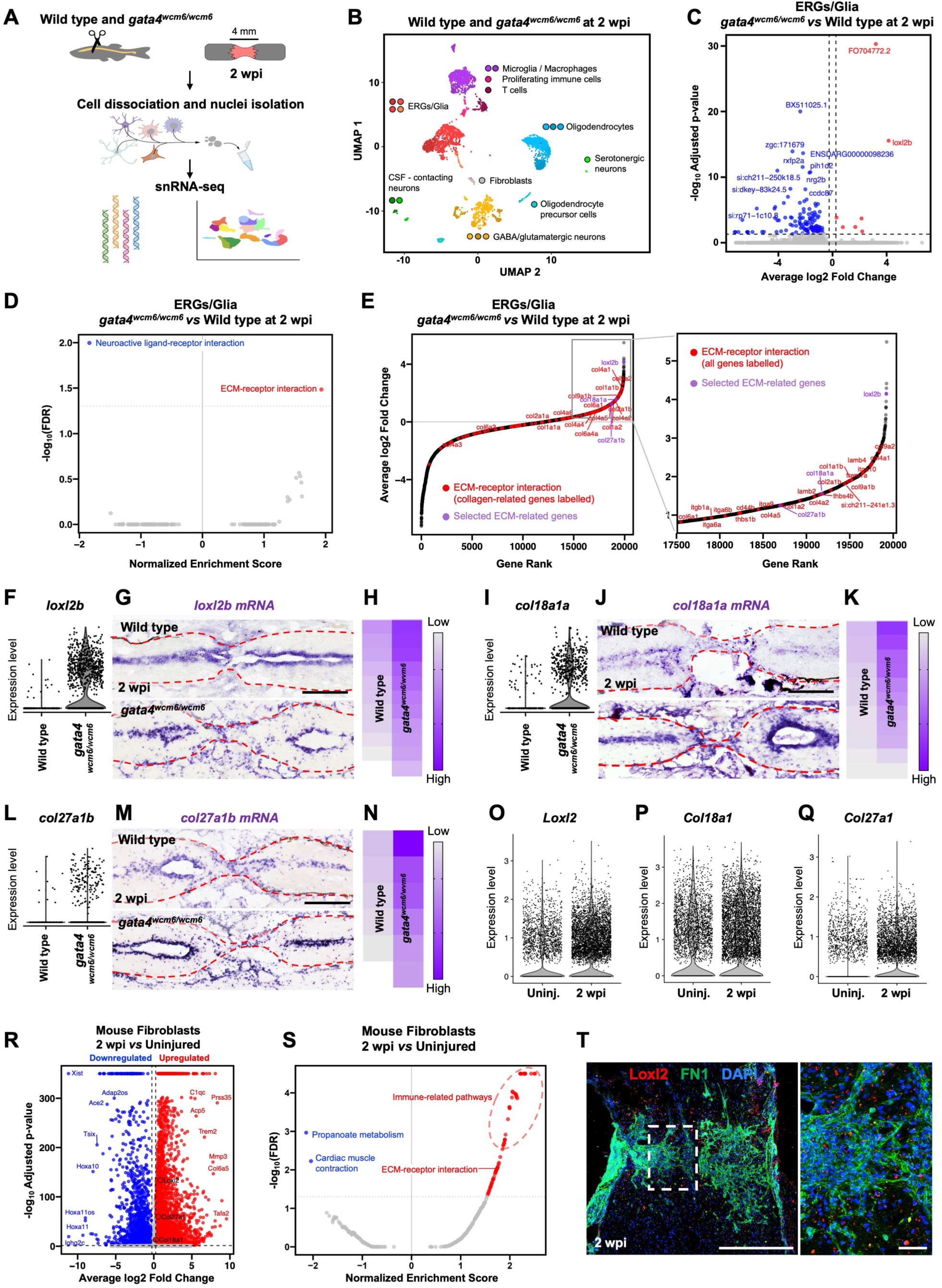
Gata4 loss-of-function induces expression of ECM-related genes. **(A)** Diagram showing single nucleus RNA-sequencing (snRNA-seq) strategy. **(B)** UMAP showing clustering of snRNA-Seq data at 2 weeks post-injury (wpi). Unbiased clustering identifies all expected cell populations, with contributions from both *gata4^wcm6/wcm6^* and wild-type cells. **(C)** Volcano plot showing differentially expressed genes in ERGs/Glia (excluding oligodendrocytes and microglia) of *gata4^wcm6/wcm6^* and wild-type spinal cords at 2 wpi. **(D)** Gene set enrichment analysis (GSEA) using the KEGG pathways database for zebrafish highlighting an extracellular matrix (ECM)-related signature in *gata4^wcm6/wcm6^* ERGs/Glia compared to wild type at 2 wpi. **(E)** Annotated Rank Plot highlighting ECM pathway genes based on log2 fold change, showing enrichment in ECM-related genes in *gata4^wcm6/wcm6^* ERGs/Glia compared to wild-type. ECM genes with the highest fold changes are highlighted in the zoomed-in inset. **(F)** Violin plot showing absolute expression levels of *loxl2b* transcripts in *gata4^wcm6/wcm6^* and wild-type ERGs/Glia at 2 wpi. **(G)** In situ hybridization for *loxl2*b on longitudinal spinal cord sections of *gata4^wcm6/wcm6^* and wild-type zebrafish at 2 wpi. Quantification shown in **(H)**. **(I)** Violin plot showing absolute expression levels of *col18a1a* transcripts in *gata4^wcm6/wcm6^* and wild-type ERGs/Glia at 2 wpi. **(J)** In situ hybridization for *col18a1a* on longitudinal spinal cord sections of *gata4^wcm6/wcm6^* and wild-type zebrafish at 2 wpi. Quantification shown in **(K)**. **(L)** Violin plot showing absolute expression levels of *col27a1b* transcripts in *gata4^wcm6/wcm6^* and wild-type ERGs/Glia at 2 wpi. **(M)** In situ hybridization for *col27a1b* on longitudinal spinal cord sections of *gata4^wcm6/wcm6^* and wild-type zebrafish at 2-wpi. Quantification shown in **(N)**. **(O-Q)** Violin plots showing upregulation of *Loxl2b*, *Col18a1a* and *Col27a1* transcripts in mouse fibroblasts at 2 wpi versus uninjured controls. **(R)** Volcano plot showing all statistically significant genes enriched in mouse fibroblasts after spinal cord injury *vs.* non injured fibroblasts at 2 wpi. **(S)** GSEA showing ECM-receptor pathway upregulation in mouse fibroblasts at 2 wpi compared to control fibroblasts. **(T)** Immunofluorescence on mouse spinal cord sections at 2 wpi showing expression of secreted Loxl2 protein (red) in and around cells expressing Fibronectin (green). Dashed lines in **G**, **J** and **M** delineate spinal cord tissue. Dashed area in **T** indicate magnified region. Scale bar in 100 μm in **G**, **J** and **L**, 500 μm in **T**, 50 μm in **T** magnified region.

### Molecular comparisons between zebrafish *gata4*-expressing glia and mammalian reactive astrocytes and fibroblasts

The enrichment of collagen and other ECM-related genes in *gata4^wcm6/wcm6^* glia led us to speculate that these cells may acquire a transcriptional program similar to mouse reactive astrocytes. To test this, we first generated a list of zebrafish genes enriched in *gata4^wcm6/wcm6^* versus wild-type glia at 2 wpi and mapped them to their mouse orthologs. This ortholog gene set was then used to examine expression in a previously published mouse reactive-astrocyte dataset^35^ **(Figure S6A)**. Comparison of differentially expressed genes (DEG) between *gata4^wcm6/wcm6^* and wild-type glia to those generated from mouse reactive versus homeostatic astrocytes revealed minimal correlation in fold-change values. GSEA of concordant and non-concordant genes did not identify shared pathways (**Figures S6B and S6C, Tables S3 and S4)**. In addition, *Loxl2, Col18a1* and *Col27a1b*, three genes we confirmed were upregulated in *gata4^wcm6/wcm6^* versus wild-type glia **(Figures 3F-N)**, were only sparsely expressed in mouse reactive astrocytes, suggesting low overall transcript abundance **(Figures S6D-F)**. These findings suggest that *gata4^wcm6/wcm6^* glia engage molecular programs distinct from those of mammalian reactive astrocytes.

Given the strong induction of *loxl2b*, *col18a1a,* and *vimentin*, genes commonly expressed in fibroblasts^36–41^, in *gata4^wcm6/wcm6^* glia, we asked whether *gata4*-mutant glia may acquire a fibroblast-like state rather than a reactive astrocyte–like phenotype following SCI. We mapped DEGs from *gata4^wcm6/wcm6^* versus wild-type glia to their mouse orthologs and assessed their expression in mouse fibroblasts after SCI compared with uninjured fibroblasts. For this analysis, we reprocessed a previously published single-cell RNA-sequencing dataset^42^, re-annotating cell types to recover fibroblast populations at 0 and 2 wpi **(Figure S7A)**. Differential expression and pathway analyses revealed extensive transcriptional remodeling of mouse fibroblasts after injury. *Loxl2b*, *Col18a1*, *Col27a1b* and additional genes including *Vim*, *Fn1*, *Col1a1* and *Col16a1*, all robustly induced in *gata4^wcm6/wcm6^* glia, were significantly upregulated in post-injury mouse fibroblasts **(Figures 3O-Q and S7A, Table S5)**. Moreover, post-injury mouse fibroblasts showed strong enrichment of immune-related pathways and upregulation of ECM–receptor interaction and collagen biosynthesis pathways **(Figures 3R, 3S and S7B, Table S6)**. These same ECM–related pathways were among the most enriched in *gata4^wcm6/wcm6^* glia **(Figure 3D and Table S2)**. Consistent with these transcriptional analyses, immunofluorescence using a secreted-LOXL2-specific antibody^43^ revealed LOXL2 at the lesion site in the adult mouse spinal cord, within the area of the fibrotic scar where fibronectin is also present **(Figure 3T)**.

Altogether, these results suggest that *gata4* loss-of-function in zebrafish redirects glia transcriptional programs toward a mouse fibroblast-like, ECM-remodeling state, rather than a canonical reactive-astrocyte response. This shift likely reflects a transition toward a fibrotic-like repair mode, highlighting a role for zebrafish *gata4* in constraining excessive matrix production during spinal cord regeneration.

### Gata4 loss-of-function increases secreted Loxl2b levels, leading to spinal cord fibrosis

*loxl2b*, one of several zebrafish paralogs within the lysyl oxidase family, was the most strongly upregulated transcript in *gata4^wcm6/wcm6^* glia compared to wild type after SCI. This protein catalyzes collagen cross-linking by converting lysine and hydroxylysine residues into reactive aldehydes (Lys^Ald^), which then form covalent bonds between collagen fibrils, creating a stable and rigid extracellular matrix **(Figures 4A and S8B)**^44^. To confirm that *loxl2b* transcriptional signal corresponds to protein-level changes, we performed immunofluorescence using an antibody validated as recognizing only the secretory mature form of Loxl2 protein^43^. This revealed a trend toward increased Loxl2 accumulation in and around the regenerating site of *gata4^wcm6/wcm6^* animals compared to wild type controls at 2 wpi, consistent with enhanced collagen remodeling **(Figures 4B, 4C and S8A)**. To assess the dynamics of collagen oxidation and cross-linking at the spinal cord lesion site, we used the oxyamine-bearing probe TMR-O, a high-precision molecular imaging probe that binds Lys^Ald^ residues generated by lysyl oxidase enzymes on collagen that is actively undergoing cross-linking^45^ **(Figure S8B).** TMR-O was previously validated for detecting Lys^Ald^ in models of fibrosis, including in zebrafish and murine hearts^45,46^. Its rhodamine moiety makes it possible to easily monitor lysyl oxidase activity on tissue sections taken from animals receiving TMR-O by systemic delivery. We first confirmed TMR-O specificity in zebrafish spinal cords by intraperitoneal injection of TMR-O or a non-binding fluorescent control probe (TMR-NB) in wild-type uninjured and 1 wpi fish. Spinal cords analyzed 4 hours post-injection showed robust TMR-O labeling around the lesion, whereas uninjured spinal cords or cords injected with TMR-NB showed no signal **(Figures S8C and S8D)**. We then administered TMR-O at 1 or 2 wpi to *gata4^wcm6/wcm6^* and wild-type animals and analyzed ECM surrounding the regenerating tissue. TMR-O fluorescence intensity was increased in mutants compared to wild types at both 1 and 2 wpi, with an approximately 70% increase at 1 wpi and a near-doubling at 2 wpi, indicating markedly enhanced formation of mature collagen cross-links during the regenerative period **(Figures 4D-F)**.

**Figure 4.**
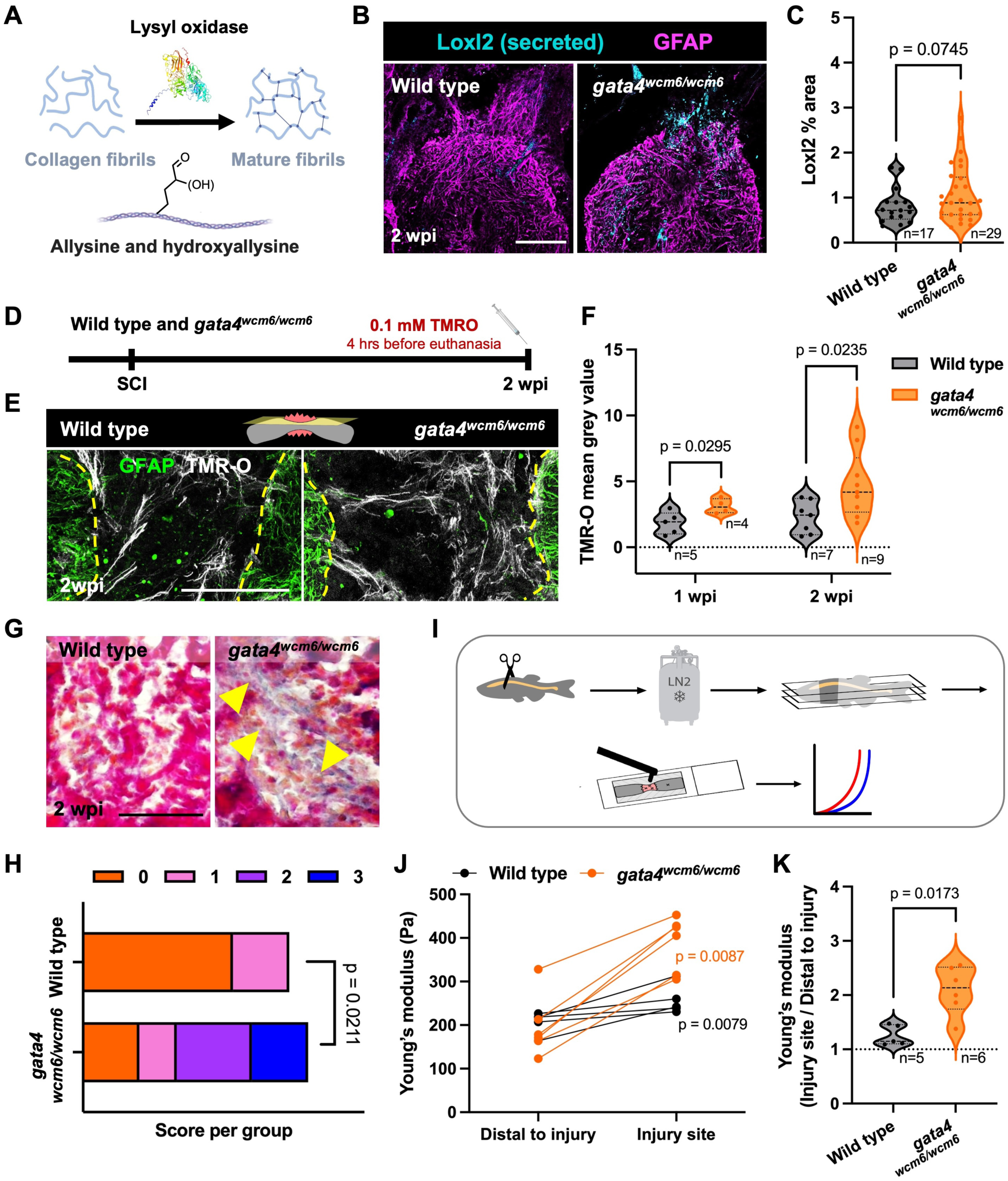
Gata4 loss-of-function promotes scarring and increases spinal cord stiffness. **(A)** Diagram of the lysyl oxidase reaction. Lysyl oxidase catalyzes the oxidation of lysine and hydroxylysine residues in collagen within the wound, generating highly reactive aldehydes (allysine and hydroxyallysine, collectively LysAld) that drive collagen cross-linking and mature fibrils formation. **(B)** Immunofluorescence staining showing increase in Loxl2 protein in wild-type versus *gata4^wcm6/wcm6^* zebrafish spinal cord at 2 weeks post-injury (wpi). Only one injury stump is shown. Quantification in **(C)**. **(D)** Experimental design for TMR-O probe administration. Animals underwent spinal cord injury and received TMR-O systemically 4 hours before tissue analyses. **(E)** Longitudinal sections of *gata4^wcm6/wcm6^* and wild-type zebrafish spinal cords stained for GFAP (green) and TMR-O fluorescence (gray) showing increased TMR-O signal in the mutant spinal cord and surrounding tissue compared to wild type. **(F)** Violin plots showing increase in TMR-O fluorescence in *gata4^wcm6/wcm6^* versus wild-type zebrafish at 1 and 2 wpi. **(G)** AFOG-stained histology sections showing increased collagen deposition in *gata4^wcm6/wcm6^* spinal cord compared to wild types. **(H)** Semiquantitative assessment of collagen deposition: 0 = no detectable aniline blue staining (no collagen deposition), 1 = low collagen deposition, 2 = moderate collagen deposition, and 3 = high collagen deposition. n = 12 *gata4^wcm6/wcm6^* and 11 wild-type animals. **(I)** Schematics of atomic force microscopy pipeline. **(J)** Comparison of the apparent Young’s moduli of *gata4^wcm6/wcm6^* and wild-type zebrafish spinal cords at 2 wpi showing differences between regions distal to the injury site and at the injury site, with higher values consistently measured into the lesion in all animals. **(K)** Ratio between Young’s moduli measured at the injury site and in regions distal to the injury, showing higher stiffness in *gata4^wcm6/wcm6^* than wild-type spinal cords at 2 wpi. Scale bar 50 μm in **B** and **G**, 100 μm in **E**. Mann-Whitney used for comparisons in **J, K**; Welch’ *t*-test used for comparisons in **C**, **F;** Fisher’s exact test used for comparisons in **H**.

To correlate TMR-O signal with fibrotic tissue deposition, we stained spinal cord sections with Acid Fuchsin-Orange G (AFOG) at 2 wpi, when TMR-O reactivity was highest. AFOG staining revealed no difference between uninjured spinal cords from wild-type or *gata4^wcm6/wcm6^* zebrafish **(Figure S9A)**. By contrast, injured *gata4^wcm6/wcm6^* spinal cords exhibited more extensive scarring than injured wild-type spinal cords **(Figures 4G and 4H)**. To assess the mechanical properties of the regenerating tissue, we performed atomic-force microscopy (AFM) indentation on non-fixed, longitudinal spinal cord sections encompassing the lesion site at 2 wpi, using a spherical probe **(Figures 4I and S8E)**. Force-distance curves were fitted with the Hertz model to derive apparent Young’s moduli (see Methods). As expected, tissue near the lesion displayed significantly increased stiffness compared to distal tissue in both *gata4^wcm6/wcm6^* and wild-type spinal cords **(Figures S8F, S8G and 4J)**^47^. Distal regions showed similar stiffness between genotypes (wild-type: 206.9 ± 24.9 Pa; *gata4^wcm6/wcm6^*: 196.2 ± 70.7 Pa), whereas lesion sites were substantially stiffer in mutants compared with wild-type animals (wild-type: 256.9 ± 33.2 Pa; *gata4^wcm6/wcm6^*: 388.4 ± 62.7 Pa) **(Figures 4J and 4K)**, consistent with increased fibrosis and collagen cross-linking. These findings show that Gata4 loss-of-function induces a pro-fibrotic state in zebrafish spinal cord, with elevated Loxl2b, increased collagen cross-linking, and stiffer ECM, which likely underlies the impaired regeneration in *gata4^wcm6/wcm6^* animals. This phenotype somewhat mirrors the fibrotic scarring observed in mouse SCI **(Figure S9B)**.

### Long-term Gata4 loss-of-function alters spinal curvature and decreases synaptic vesicles

To investigate why functional deficits persist in *gata4^wcm6/wcm6^* at 4 wpi despite histological evidence of (albeit delayed) substantial glial and axonal regeneration **(Figures 2E-M)**, we analyzed older *gata4^wcm6/wcm6^* zebrafish. Since spinal curvature defects are frequently observed during and after regeneration and can affect swimming performance, we examined post-injury scoliosis in mutants and wild types at 2, 4, and 6 wpi. For both *gata4^wcm6/wcm6^* and wild type fish, no significant differences were detected before injury or up to 4 wpi, but by 6 wpi, *gata4^wcm6/wcm6^* animals developed pronounced spinal curvature (p = 0.0157, Mann-Whitney) **(Figures S10A and S10B)**. This delayed scoliosis likely reflects persistent, progressive and intransigent fibrotic remodeling and increased tissue stiffness in absence of Gata4 function, potentially resulting from collagen cross-linking and matrix deposition in mutants. To determine whether these structural alterations also affected neuronal function, we assessed synaptic integrity. Analysis revealed a significant reduction in synaptic vesicles in *gata4^wcm6/wcm6^* spinal cords, as indicated by decreased immunoreactivity for the presynaptic markers Syt2a at 4 wpi and SV2 at 6 wpi, compared to wild-type fish **(Figures S10C-F)**. Collectively, these results indicate that chronic Gata4 loss-of-function impairs both structural and functional aspects of regeneration after SCI.

### Inhibition of Loxl2b rescues regeneration defects in *gata4^wcm6/wcm6^* mutants

To determine if the phenotype observed in *gata4^wcm6/wcm6^* zebrafish could be ameliorated by blocking the elevated Loxl2b activity, we treated animals with β-aminopropionitrile (BAPN). This potent and irreversible inhibitor of lysyl oxidases has been extensively characterized and used across multiple model organisms, including zebrafish^48^. We opted for broad pharmacological inhibition rather than cell-specific genetic ablation to counteract potential contributions from Loxl2b secreted by non-glial cell types at the injury site **(Figure 1A)**. *gata4^wcm6/wcm6^* animals were injured and incubated with BAPN for two weeks **(Figure 5A)**. BAPN administration reversed collagen deposition in and around the lesion observed in *gata4^wcm6/wcm6^* animals, reducing it to levels comparable to wild type, as assessed by AFOG staining **(Figures 5B and 5C)**. We also analyzed 2 wpi spinal cords from zebrafish treated with BAPN and injected with the TMR-O probe. BAPN treatment reduced TMR-O labeling in *gata4^wcm6/wcm6^* lesions to levels comparable with those of regenerating wild-type animals **(Figures 5D and 5E)**. We assessed functional recovery following injury by subjecting wild-type, *gata4^wcm6/wcm6^*, and BAPN-treated *gata4^wcm6/wcm6^* animals to swim tests. Uninjured animals showed no differences in swim capacity across groups (wild-type, *gata4^wcm6/wcm6^*, and BAPN-treated *gata4^wcm6/wcm6^*) **(Figure 5F)**. Following SCI, *gata4^wcm6/wcm6^* animals consistently exhibited a trend toward reduced swim recovery compared with wild-type animals, although this difference did not reach statistical significance by Mann-Whitney test, likely due to inter-individual variability. Notably, BAPN treatment restored swim performance in *gata4^wcm6/wcm6^* animals to levels comparable with wild type, offsetting the reduced recovery observed in untreated mutants **(Figure 5G)**. These results correlated with histological analyses of spinal cord tissue, where BAPN-untreated *gata4^wcm6/wcm6^* animals showed a 38% and 47% reduction in glial and axon bridging, respectively, compared to controls. Remarkably, BAPN treatment completely rescued these defects, restoring glial and axon bridge formation to wild-type levels **(Figure 5H-J)**. Together, these results indicate that BAPN treatment rescues excessive collagen deposition, glial and axonal bridging and, at least partially, the functional recovery defect in *gata4^wcm6/wcm6^* animals following injury.

**Figure 5.**
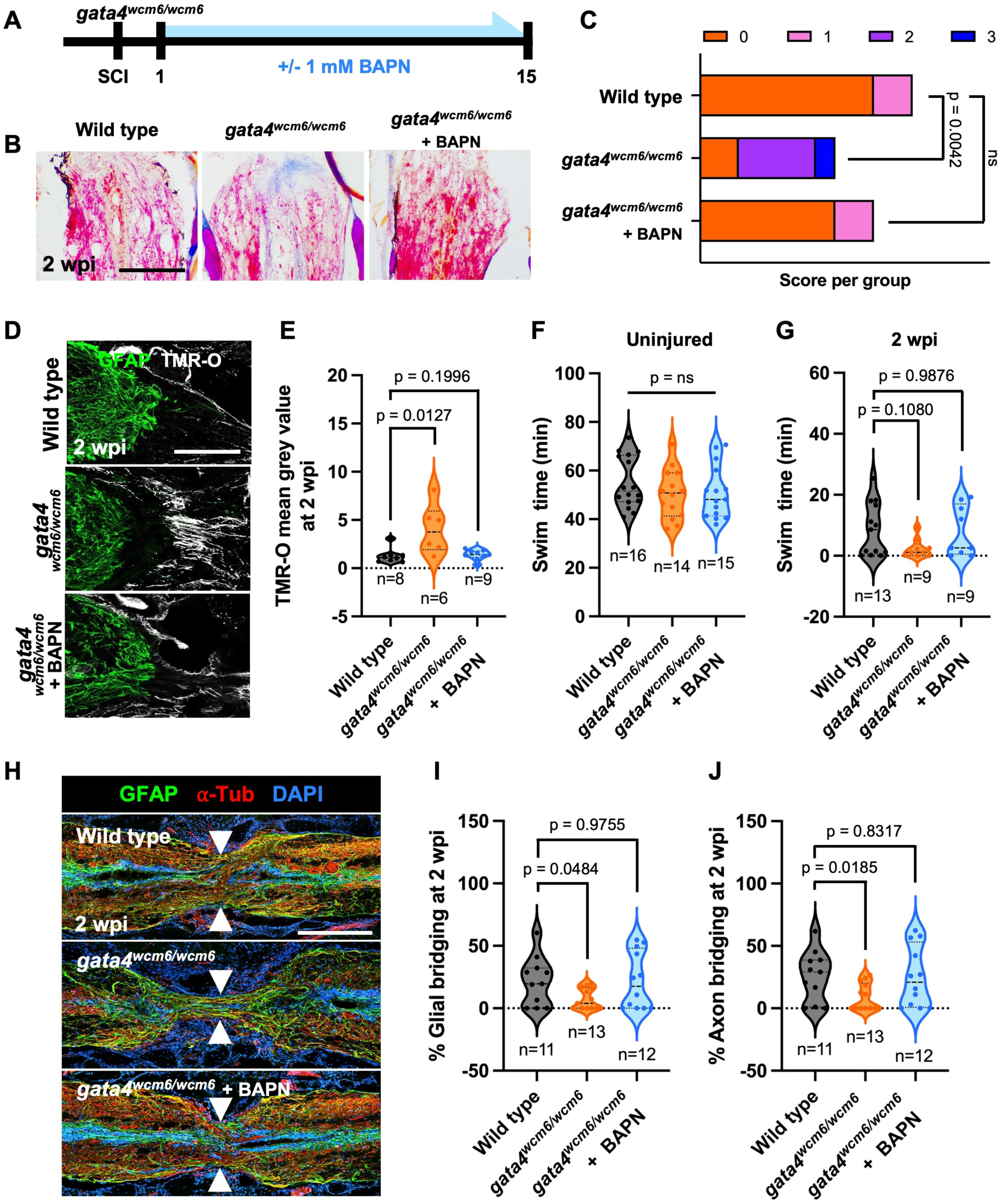
Pharmacological inhibition of Loxl2 reduces fibrosis and restores tissue bridging and function in *gata4^wcm6/wcm6^* spinal cords. **(A)** Schematic showing BAPN administration protocol. **(B)** AFOG-stained histological sections showing increased collagen deposition in the spinal cord of *gata4^wcm6/wcm6^* mutants compared to wild-type controls, and rescue of this fibrotic response in *gata4^wcm6/wcm6^* treated with BAPN at 2 weeks post-injury (wpi). **(C)** Semiquantitative assessment of collagen deposition: 0 = no detectable aniline blue staining (no collagen deposition), 1 = low collagen deposition, 2 = moderate collagen deposition, and 3 = high collagen deposition. n = 11 wild type, 7 *gata4^wcm6/wcm6^* and 9 *gata4^wcm6/wcm6^* +BAPN animals. **(D)** Longitudinal sections of wild-type, *gata4^wcm6/wcm6^*, BAPN-treated *gata4^wcm6/wcm6^* spinal cords stained for GFAP (green) and TMR-O fluorescence (gray), showing that BAPN treatment rescues the *gata4^wcm6/wcm6^* phenotype to wild-type levels. Quantification of TMR-O signal is shown in the violin plots in panel **(E). (F)** Endurance swim testing of uninjured wild-type, *gata4^wcm6/wcm6^* and BAPN-treated *gata4^wcm6/wcm6^* zebrafish. Animals were placed in an enclosed swim tunnel with increasing water current, and the time to exhaustion was recorded. **(G)** Endurance swim testing of injured wild-type, gata4^wcm6/wcm6^ and BAPN-treated gata4^wcm6/wcm6^ zebrafish. Swimming capacity against progressively increasing currents was measured at 2, 4, and 6 wpi. **(H)** Spinal cord tissue sections of wild-type, *gata4^wcm6/wcm6^* and *gata4^wcm6/wcm6^* + BAPN zebrafish at 2 wpi, stained for the glial marker GFAP (green) and axonal marker acetylated α-Tubulin (red). Arrows indicate sites of tissue bridging. Quantification of glial bridge and axon bridge shown in **(I)** and **(J),** respectively. Scale bar 100 μm in **B** and **D**, 150 μm in **H**. Fisher’s exact test used for comparisons in **C**. Mann-Whitney used for comparisons in **E**, **F, G**, **I**, **J.**

### Hif1α contributes to the post-SCI induction of *gata4* via an upstream cis-regulatory region

To gain insight into the signals that activate *gata4* after SCI, we performed a first-pass pharmacological screen in zebrafish larvae to test whether injury-associated cues can de-repress *gata4* expression. We leveraged *Tg(−14.8kb-gata4:EGFP; gfap-H2A:mCherry)* larvae, which regenerate the spinal cord more rapidly than adults and whose optical transparency and small size enable direct, longitudinal in vivo visualization of GFAP⁺, mCherry-labeled glia and *gata4*:EGFP reporter activity in the same animals over time^49,50^. Tissue injury triggers a cascade of events, including mechanical stress, hypoxia, reactive oxygen species (ROS) production, inflammation, and apoptosis **(Figure 6A)**. Uninjured larvae at 2 days post fertilization (dpf) were exposed to agents mimicking these injury-associated cues, and *gata4*:EGFP expression in the spinal cord was monitored over time. Agents we used included Dimethyloxalylglycine (DMOG), a pan-hydroxylase inhibitor that stabilizes the Hif1α protein and activates hypoxic signaling^51,52^; phenylhydrazine hydrochloride (PHZ), which induces anemia through red blood cell lysis^53,54^; and hydrogen peroxide (H₂O₂), previously shown to stimulate ROS generation^55^ **(Figure 6B)**. Using uninjured larvae allowed us to identify candidate upstream activators of *gata4* expression while avoiding confounding effects of the injury itself. Among these treatments, constant exposure to DMOG induced robust *gata4:EGFP* expression in spinal cord glia at 5 dpf. This indicates that hypoxic signaling is sufficient to at least partially activate *gata4* transcription in the absence of injury **(Figures 6C and 6D, Movie S1)**.

**Figure 6.**
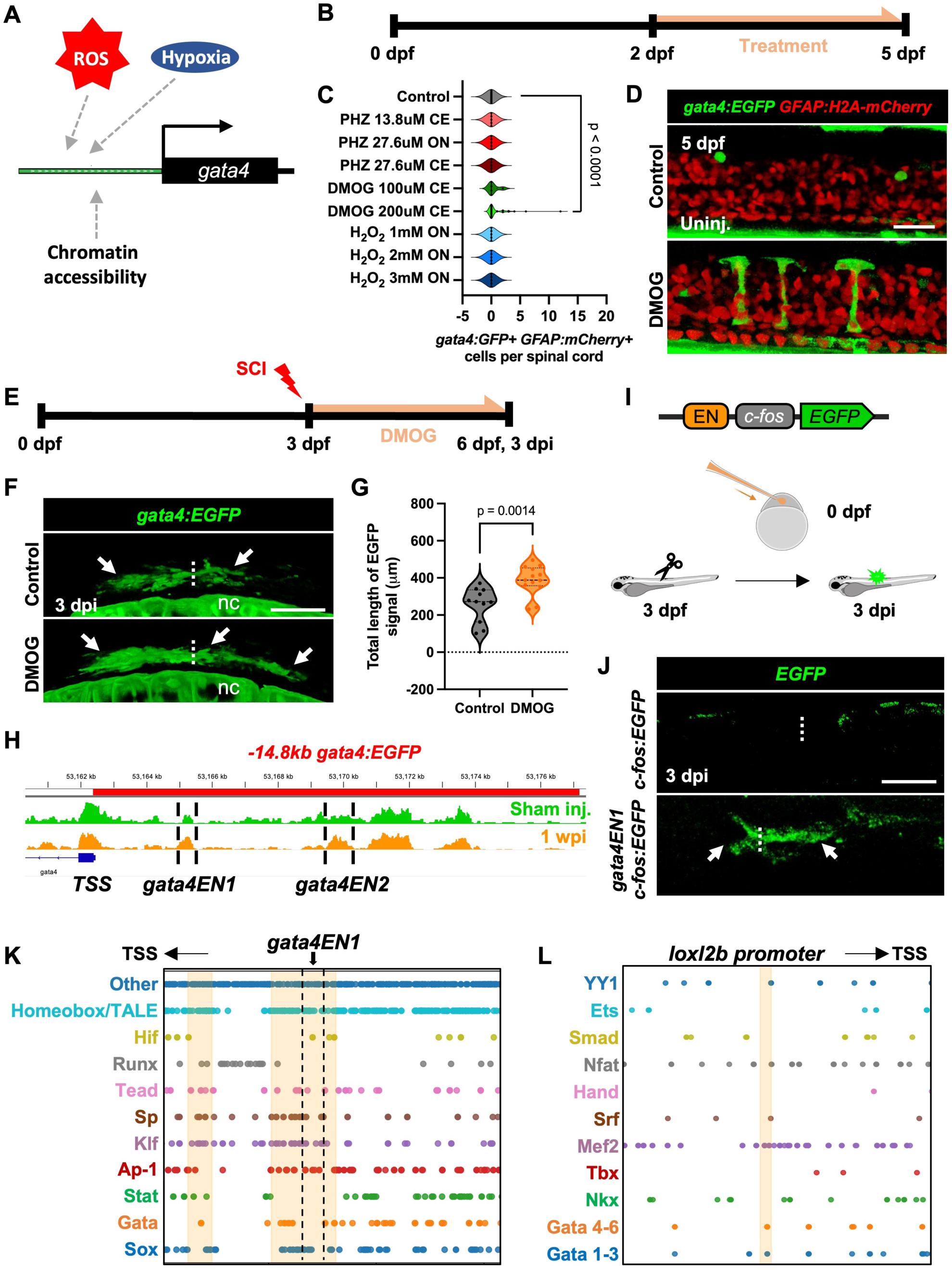
Hif1α contributes to injury-induced *gata4* expression through an upstream cis-regulatory region. **(A)** Schematic showing selected injury-induced signals acting as possible inducers of *gata4* expression. **(B)** Experimental design for larval drug and compound treatments in uninjured animals. **(C)** Absolute number of *gata4:EGFP⁺* glial cells in *Tg(−14.8kb-gata4:EGFP; gfap:H2A-mCherry)* uninjured larvae treated with phenylhydrazine (PHZ), DMOG, or H₂O₂. ON, overnight; CE, constant exposure. **(D)** Representative images of *Tg(−14.8kb-gata4:EGFP; gfap:H2A-mCherry)* larval spinal cords after DMOG exposure (200 mM). GFAP labels glial nuclei. **(E)** Experimental design for larval DMOG treatment following spinal cord transection injury. **(F)** Representative images of *Tg(−14.8 kb gata4:EGFP)* larvae at 1 day after DMOG treatment, showing increased *gata4:EGFP* expression in treated animals compared with controls. nc indicates the notochord, previously shown to activate *−14.8kb-gata4:EGFP* during development. **(G)** Quantification of *−14.8kb-gata4:EGFP* signal in the spinal cord. **(H)** ATAC-seq genome browser tracks showing two regions of increased chromatin accessibility in the *−14.8kb gata4* upstream regulatory region at 1 week post injury compared with uninjured controls (*gata4EN1* and *gata4EN2*). **(I)** Strategy used to assess *in vivo gata4EN1* and *gata4EN2* enhancer function. **(J)** Immunofluorescence images showing injury-induced expression driven by *gata4EN1* at 2 days post-injury. **(K)** FIMO motif analysis of the *-14.8kb gata4* upstream regulatory region. Predicted transcription factor binding sites are shown by family and position relative to the transcription start site. The shaded region marks 2 motif-dense regions identified using defined clustering criteria (>5 TF families, >8 motif hits, q < 0.05), illustrating a modular upstream regulatory region architecture that includes predicted transcription factor binding sites. **(L)** FIMO motif analysis of the *loxl2* upstream regulatory region showing predicted transcription factor binding sites shown by family and position relative to the transcription start site. The shaded region marks a motif-dense region identified using the same clustering criteria as in **K**, illustrating a modular upstream regulatory region architecture that includes predicted GATA-binding sites. A selected portion of the full −14.8kb region is shown in panels **K** and **L;** the complete upstream regulatory region is shown in **Figure S11**. Scale bar 20 μm in **D**, 100 μm in **F** and **J.** Mann-Whitney and Welch’ *t*-test used for comparisons in **C** and **G**, respectively.

We next combined DMOG treatment with spinal cord transection, reasoning that DMOG exposure might enhance endogenous *gata4:EGFP* activation in an injury context. Larvae undergoing spinal cord transection at 3 dpf were continuously exposed to DMOG and imaged daily **(Figure 6E)**. EGFP production was quantified along the spinal cord, rostral and caudal to the lesion, at 3 dpi. DMOG-treated animals showed a ∼51.2% increase in the total length of EGFP signal, with similar increase rostral and caudal to the lesion site, compared to vehicle-treated controls, despite variability across individual fish. This result suggests that DMOG exposure can elevate *gata4* expression beyond the levels normally observed following injury **(Figures 6F and 6G)**.

To identify *cis*-acting regulatory DNA sequences driving localized *gata4* activation after SCI, we analyzed spinal cord chromatin accessibility datasets we previously generated^14^ focusing on the −14.8kb upstream regulatory region of the *Tg(−14.8Kb-gata4:EGFP)* reporter line. This analysis identified two regions with increased chromatin accessibility at 1 wpi compared with sham-injured controls, in which skin and muscle were cut but the spinal cord remained intact (indicated by dashed lines, **Figure 6H**). One dynamically accessible sequence, *gata4EN1*, of ∼620 bp was located about −2.7 kb relative to the *gata4* transcriptional start site (TSS) while the other, *gata4EN2,* was ∼680 bp in length at −7.3 kb upstream of the TSS. To test whether these regions can direct injury-induced gene expression after SCI, we placed them upstream of the minimal promoter *c-fos,* normally silent, and an EGFP reporter gene. Constructs were injected into fertilized zebrafish eggs, which underwent spinal cord transection injury at 3 dpf and EGFP signal was assessed at the lesion site by *in vivo* imaging at 4, 5 and 6 dpf **(Figure 6I)**. We did not detect spinal cord fluorescence in uninjured *gataEN1-cfos:EGFP* or *gataEN2-cfos:EGFP* animals, nor in animals carrying a *cfos:EGFP* control construct. Following SCI, *gataEN2-cfos:EGFP* and *cfos:EGFP* control animals showed no EGFP signal, whereas *gataEN1* directed EGFP expression at SCI sites **(Figure 6J)**. This pattern of expression closely mirrored that observed in *Tg(−14.8kb-gata4:EGFP)* larvae. Cells expressing EGFP in the spinal cord were identified as glial cells by testing for colocalization with the glial marker GFAP **(Figure S11A).**

To further understand regulation of *gata4* expression after injury, we performed motif scanning with the MEME suite FIMO function across the *-14.8kb gata4* region, selecting a broad range of TF families implicated in at least one of the following programs: regeneration, stress/inflammation, chromatin remodeling, mechanical response, development, and pioneer TFs. This analysis revealed a dense, non-random distribution of predicted TF binding sites for several major TF families **(Figures 6K and S11B, Tables S7 and S8)**. Notably, several of the top enriched motifs included Homeobox/TALE, Hif, Runx, Tead, Sp, Klf, Ap-1, Stat, Gata, and Sox. These factors are involved in key developmental and cellular processes such as lineage specification (Homeobox/TALE, Gata, Sox), hypoxia response (Hif), proliferation and differentiation (Runx, Tead, Sp, Klf), stress and signaling responses (Ap-1, Stat), and transcriptional regulation during organogenesis. Two prominent TF motif clusters reached our selected threshold (red-shaded in **Figure 6K**): one located approximately ∼0.5-1.1 kb upstream of the TSS and a second at ∼2-3 kb upstream of the TSS that, importantly, overlaps with *gata4EN1*. Analyses of TF motif clusters located ∼2-3 kb from the TSS, containing *gata4EN1,* suggests that *gata4* transcription is regulated by synergistic interactions between localized regulatory hotspots. Hif1A motifs, along with Klf and Sp motifs showed the highest significance based on p- and q-values **(Figure 6K)**^56^. Notably, Klf and Sp TFs physically and functionally interact with Gata4, often forming complexes to cooperatively regulate gene expression^57,58^. These interactions are particularly important in contexts such as heart development and cancer, where they influence cell proliferation, differentiation, and cardiac hypertrophy through shared binding sites and regulatory crosstalk. Together, these findings suggest that after SCI in zebrafish Hif1α activates *gata4* either directly binding within the upstream regulatory region, or indirectly via activation of genes coding for Klf and Sp factors, that subsequently bind *gata4* cis-regulatory regions. It is possible that Hif1α-mediated direct and indirect activation both contribute, with the latter supported by previous studies demonstrating Hif1α regulation of Sp and Klf in other contexts^57,58^.

Finally, to investigate mechanisms by which Gata4 regulate *loxl2b* transcription, we analyzed the *loxl2b* upstream regulatory region for potential TF sites using FIMO motif scanning. Mapping motif occurrences across the −5kb region upstream of the TSS revealed a non-uniform distribution, again with discrete regions containing a relatively high density and diversity of TF motifs **(Figures 6L, S11B and S11C, Tables S9 and S10)**. Applying clustering criteria scanning the −5kb *loxl2b* upstream regulatory region by narrow 75 bp windows containing motifs from more than five TF families, more than eight total motif hits, and at least one motif with low FDR (*q* < 0.05), focused us on a prominent motif-dense region ∼3.8kb upstream of the *loxl2b* TSS. This cluster harbors consensus motifs for multiple TF families, including Gata, Nkx, Mef2, Srf, and Ets, consistent with a potential *cis*-regulatory module. The number of GATA motifs within this region supports the possibility that Gata4 directly contributes to *loxl2b* repression, potentially in co-operation with other TFs such as Mef2 which has been previously described to co-bind with Gata4 to regulate gene expression through upstream regulatory region binding^59^. Together, these analyses indicate that *loxl2* is embedded within a complex regulatory landscape and yield a mechanistic model in which Gata4 influences ECM remodeling via transcriptional control of the collagen cross-linking enzyme Loxl2b.

## DISCUSSION

Our study identifies Gata4 as a central regulator that preserves glial identity and pro-regenerative competence after SCI in zebrafish. *gata4* is expressed primarily in a subset of ERGs proximal to the rostral and caudal spinal cord stumps and in glial processes extending across the lesion site after SCI, and its loss of function unleashes a transcriptional, and possibly phenotypic, shift toward a fibrotic, anti-regenerative state. Gata4-deficient glia, and possibly additional fibroblast-like cells in and around the lesion, upregulate ECM components and cross-linking enzymes, leading to excessive collagen deposition, tissue stiffening, and impaired neural repair. In contrast to the transient, permissive ECM normally produced during zebrafish spinal cord regeneration, the Gata4-deficient lesion resembles in several ways the persistent fibrotic scar formed after SCI in mammals. These findings establish that active suppression of fibrotic gene-expression programs is a prerequisite for regenerative repair and position Gata4 as a transcriptional safeguard that prevents maladaptive matrix remodeling.

The ECM components that accumulate in Gata4-deficient lesions are the same that dominate the mammalian fibrotic scar, where they increase mechanical stiffness, restrict cell migration, and inhibit axonal regrowth ^1^. Among these, Loxl2b emerged as a key effector: its expression and enzymatic activity were markedly increased in Gata4-deficient glia, driving enhanced collagen cross-linking and matrix stabilization. Pharmacological inhibition of Loxl2b with BAPN reduced matrix rigidity and restored glial bridge–forming capacity in Gata4-deficient zebrafish, underscoring the functional importance of Gata4-dependent control of ECM cross-linking. Given that GATA4 also suppresses fibrogenic programs in heart and liver repair in mammals^23,60^, our findings point to a conserved anti-fibrotic role for this factor across regenerative contexts and species.

We find that using DMOG as a proxy for Hif1α stabilization is sufficient to induce *gata4* expression in *uninjured* glia, suggesting that at least some hypoxia-sensing pathways are part of initiating *gata4* derepression and the pro-regenerative transcriptional cascade. We propose that localized hypoxia, likely enhanced at the injury site, activates Hif1α, that through consensus motifs in upstream *gata4 cis*-regulatory domains elicits a transcriptional program that suppresses fibrotic remodeling and sustains glial plasticity. GATA4’s role in suppressing Loxl2b-mediated cross-linking maintains a compliant ECM that supports appropriate cell migration and axonal extension, facilitating the rebuilding of tissue architecture, neuronal connectivity, and function. Identifying transcription factors that link environmental stress signals to tissue mechanics, and that ensure glial activation promotes regeneration rather than fibrosis, is likely critical for understanding the limited regenerative capacity in mammals, including humans.

Using DMOG to stabilize Hif1α induced *gata4* expression in uninjured glia, whereas phenylhydrazine (PHZ)-induced anemia had no detectable effect. Although both compounds were delivered by bath immersion and could act broadly, their mechanisms likely influence the spatial pattern of hypoxia signaling. DMOG directly stabilizes Hif1α at the cellular level, preferentially amplifying hypoxia-responsive transcription in cells already primed for this pathway. In contrast, PHZ reduces systemic oxygen-carrying capacity but may not generate sufficient or sustained local hypoxia within the spinal cord to activate *gata4*. Without direct measurements, it remains unclear whether PHZ treatment produced substantial tissue-level hypoxia. Together, these findings indicate that local or cell-autonomous Hif1α activation, rather than global oxygen deprivation, serves as a key signal initiating *gata4* derepression and the pro-regenerative transcriptional program.

The Gata4 upstream regulatory region contains discrete clusters of various TF motifs binding sites, suggesting a convergent, highly interlinked architecture that incorporate numerous upstream inputs on multiple transcriptional regulators. HIF1A motifs are present in these clusters, and KLF and SP sites also appear prominently. Both factors have been reported as modulated by hypoxia-induced HIF1A, plausibly providing an indirect but still rapid mechanism for *gata4* transcriptional de-repression. Having direct and indirect regulatory programs working together could provide for flexible and buffered tuning according to the different responses appropriate to and relevant for the specific biological context.

These findings go beyond previous concepts for the action of glial cells in spinal cord regeneration. Beyond forming a passive structural bridge, in adult zebrafish glial cells actively prevent fibrotic scar formation and maintain a permissive environment for axonal regrowth. By coordinating ECM remodeling and glial state transitions, Gata4 enables the zebrafish spinal cord to heal without forming a persistent scar. Translationally, our data suggest that enhancing GATA4 activity or inhibiting collagen cross-linking enzymes such as LOXL2, through agents like BAPN, could mitigate fibrosis and promote repair in the injured mammalian CNS. Notably, a 2001 study reported that BAPN-mediated lysyl oxidase inhibition significantly improved motor recovery after SCI in mice^61^ but the underlying cellular mechanisms – particularly those related to axon regeneration as seen in our zebrafish model – remain largely unexplored. Furthermore, we consider the zebrafish *gata4^wcm6/wcm6^* model, where other pro-regenerative processes work synergistically to repair tissue structure and function, to be a valuable platform to test anti-fibrotic compounds and uncover new regulators of scar formation.

While our findings identify Gata4 as a key regulator of ECM composition during spinal cord regeneration, several important questions remain. DMOG, which stabilizes Hif1α in the presence of oxygen, also inhibits certain histone demethylases, so some effects on *gata4* activation may not be purely Hif1α-mediated. Future ChIP-seq analyses will be required to determine whether Gata4 directly regulates *loxl2* expression, and to dissect direct versus indirect effects of Hif1α on *gata4* regulatory regions. The *gata4^wcm6/wcm6^* mutation is global, affecting glia, fibroblasts, and other cell types, limiting assignment of cell-type–specific roles. Finally, although tissue bridging recovers in mutants, swimming deficits persist. We assessed only swim endurance in a closed tunnel, so more subtle impairments in fine motor control or swim kinematics, such as tail beat and stride length, remain undetected. Finer circuit-level analyses will be required to fully understand the persistence of these functional impairments.

Together, these insights establish GATA4 as a key transcriptional node linking hypoxic signaling, glial plasticity, and ECM dynamics, and highlight strategies to harness regenerative programs across species. Dissecting the molecular and cellular mechanisms underlying zebrafish spinal cord regeneration may facilitate reverse engineering these processes to inform regenerative strategies in mammals. This comparative, cross-species approach positions zebrafish not only as a model for basic biology but also as a guide for translational strategies aimed at overcoming the fibrotic barrier that limits spinal cord regeneration in humans.

## MATERIAL AND METHODS

### Zebrafish

Wild-type, mutant, or transgenic zebrafish of the Ekkwill (EK) strain were used for all experiments. Control and experimental animals were female and male clutch mates aged 3 to 9 months, measuring ∼2.5 cm in length. Zebrafish were raised under standard laboratory conditions at 28.5°C. Experimental fish and size-matched control siblings with balanced sex representation were used for all assays. Spinal cord transections and subsequent regeneration analyses were performed blinded to genotype when possible, and each experiment was replicated two to four times using independent clutches of animals. All experiments were conducted in accordance with guidelines established by the Institutional Animal Care and Use Committee and the Institutional Biosafety Committee at Vanderbilt University (protocol M2400047), as well as French regulations for animal care, with approval from the local Animal Care and Use Committee of the Institute of Biology Valrose (Nice) and the French Ministry of Research (no. 43195-2023060312002720). The following previously published lines were used for this study: *Tg(−14.8kb-gata4:EGFP)*^29^*, Tg(huC:dsRED)*^62^*, Tg(gfap:H2A-mcherry)*^50^*, Tg(gfap:EGFP)*^63^*, Tg(−14.8kb-gata4:dsRed2)*^64^*, Tg(olig2:EGFP)*^65^, gata4^wcm6/wcm6^ ^32^.

### Spinal cord injury

Adult zebrafish were anesthetized in 0.02% aminobenzoic-acid-ethyl methyl-ester (MS222, Sigma). Scales were removed, and a small incision was made through the dorsal musculature using fine scissors to expose the vertebral column, which was then transected midway between the dorsal fin and the operculum. Full spinal cord transection was visually confirmed at the time of surgery, and complete loss of swimming ability at 1, 2 and 3 dpi.

Zebrafish larvae at 3 dpf were anaesthetized in 0.02% aminobenzoic-acid-ethyl methyl-ester (MS222, Sigma). Larvae were transferred to a petri dish. Following removal of excess water, larvae were placed in a lateral position, and the tip of a sharp 30½ G syringe needle was used to inflict a transection injury on the dorsal part of the trunk at the level of the 15th myotome.

### Immunofluorescence

At the time of analysis, spinal cords were collected, fixed overnight in 4% paraformaldehyde, rinsed in phosphate buffer, and cryoprotected in 20% sucrose. Samples were embedded in OCT (Tissue-Tek), frozen on dry ice, and cryosectioned at 20 μm (longitudinal) or 16 μm (transverse). For standard immunofluorescence, sections were rehydrated in phosphate buffer, permeabilized in 0.2% Triton X-100, and blocked for 1 hour at room temperature in bovine serum albumin-blocking reagent. Sections were incubated overnight with primary antibodies diluted in blocking reagent, washed in phosphate buffer, and incubated for 1 hour with secondary antibodies and DAPI (Thermo Fisher Scientific, D3571, 1:5000). Following washes, sections were mounted using Fluoromount-G (Invitrogen, 00-4958-02). LOXL2 staining followed the same procedures with the addition of an antigen retrieval step. Sections underwent antigen retrieval in sodium citrate buffer at 98°C for 6 minutes and were allowed to cool for 20 minutes before proceeding with standard immunofluorescence. Primary antibodies included: rabbit anti-GFP (Life Technologies, A11122, 1:200), mouse anti-GFAP (ZIRC, Zrf1, 1:1000), rabbit anti-GFAP (Sigma, G9269, 1:200), rabbit anti-Sox2 (Abcam, ab97959, 1:200), mouse anti-HuC/D (Invitrogen, A-21271, 1:100), mouse anti-Acetylated-α-tubulin (Sigma, T6793, 1:1000), mouse anti-PCNA (Sigma, T6793, 1:500), rabbit anti-Fibronectin (Sigma-Aldrich F3648, 1:200), mouse anti-Vimentin (Proteintech 60330-1-lg, 1:500), mouse anti-Loxl2 (Proteintech, 67139-1-Ig), mouse anti-Znp-1 (DSHB, AB2315626, 1:150), mouse anti-SV2 (DSHB, AB2315387, 1:150). Secondary antibodies (Life Technologies, 1:200) were highly cross-absorbed Alexa Fluor 488/546/594 goat anti-rabbit or anti-mouse. All confocal images were acquired on Zeiss LSM 880 and Nikon AX/R with NSPARC detection microscopes.

### In Situ Hybridization

In situ hybridization was performed on cryosections of paraformaldehyde-fixed spinal cord as previously described^66^. To generate probes, gBlock fragments containing target sequences upstream of a T7 upstream regulatory region were ordered from IDT. Probes were transcribed using T7 RNA polymerase (M0251, New England BioLab). In situ hybridization signals were visualized by immunoassay using an anti-DIG-AP (alkaline phosphatase) antibody (11093274910, Sigma-Aldrich) and subsequent catalytic color reaction with NBT (nitroblue tetrazolium) (11383213001, Sigma-Aldrich) / BCIP (5-bromo-4-chloro-3-indolyl-phosphate) (11383221001, Sigma-Aldrich). Sections were imaged using an Olympus BX41TF microscope.

### AFOG staining

Fish underwent spinal cord transection and were allowed to recover for 2 weeks. At 2 wpi, animals were euthanized, and spinal cords were dissected and fixed overnight in 4% paraformaldehyde. Tissues were then washed in 0.5 M EDTA at 4 °C for 3 days, with EDTA refreshed every 1.5 days. Samples were subsequently cryoprotected in 20% sucrose for 1–3 overnights, embedded, and longitudinally sectioned at 20 µm. For AFOG staining, sections were hydrated in deionized (DI) water for 10 minutes and incubated in preheated Bouin’s Fixative Solution (1120-16, BICCA) for 2 hours at 60 °C. Slides were then allowed to incubate for an additional hour at room temperature, followed by gentle washing under constant running water for 30 minutes. Sections were incubated in 1% phosphomolybdic acid (P1910, TCI) for 5 minutes and rinsed in DI water for 5 minutes. Slides were stained for 5 minutes in AFOG solution containing 0.25 g aniline blue (GRM901-25G, HIMEDIA), 0.5 g orange G (S25453, Fisher), and 0.75 g acid fuchsin (227900250, Thermo Scientific) dissolved in 50 mL total volume. Sections were then rinsed twice in DI water for 5 minutes each, dehydrated through graded ethanol washes (75%, 95%, and 100% ethanol; two washes each for 5 minutes), cleared in xylene for 2 minutes, briefly air-dried, and mounted with mineral oil.

### Drug and probe treatments

For β-aminopropionitrile (BAPN; Cat. No. E4275, Selleckchem, Houston, TX) treatment, adult fish were allowed to recover for 1 day following SCI and then transferred to 300 mL of system water supplemented with an air stone. Experimental animals were treated with BAPN at a final concentration of 1.0 mM, and BAPN-containing water was refreshed every other day. Control *gata4^wcm6/wcm6^* and wild-type fish underwent water changes on the same schedule. Treatment was maintained under continuous exposure from 2 to 14 dpi, after which fish were euthanized, and spinal cord tissue was collected for histological analysis.

For TMR-O and TMR-NB treatments, at 2 wpi fish were injected intraperitoneally with 10 µL of 0.1 mM TMR-O or TMR-NB. Animals were kept in the dark for 4 hours following injection, then euthanized, and tissues were processed for histological analysis.

For larval drug exposure experiments, DMOG was dissolved in DMSO to a stock concentration of 2 mg/ml, and working solutions (100 or 200 μM) were prepared by dilution of the stock in fish water. Phenylhydrazine was dissolved in DMSO to a stock concentration of 2 mg/ml, and working concentrations (13.8 or 27.6 μM) was prepared by dilution in fish water. Hydrogen peroxide (H₂O₂; Fisher Scientific, H325-500) was added directly to fish water to prepare working solutions at a final concentration of 1, 2 or 3 mM. Larvae were treated in 24-well plates, with five larvae per well in a total volume of 1 ml and were subjected to either continuous drug exposure or overnight treatment.

### Swim capacity assays

To assess swim capacity, groups of fish were placed in a 5.0 L swim tunnel respirometer (Loligo, SW100605L, 120V/60Hz), as previously described^17^. Following a 20-minute acclimation period at a low, constant water current, the flow velocity was increased every two minutes. Fish were allowed to swim against the current until exhaustion or until reaching a 2 hour maximum duration. Exhausted animals were removed from the chamber without disturbing the remaining fish, while others continued to swim. For each fish that reached exhaustion, swim time in minutes and the corresponding water velocity in cm/sec were recorded and used as measurement of swim capacity. After all fish swam to exhaustion, they were allowed to recover and then placed back in recirculating water.

### Atomic force microscopy

At 2 wpi, spinal cords were dissected and incubated for 1 hour on ice in an artificial cerebrospinal fluid (aCSF)–like buffer supplemented with protease inhibitors (134 mM NaCl, 2.9 mM KCl, 1.2 mM MgCl₂, 2.1 mM CaCl₂, 10 mM HEPES, 10 mM glucose; pH 7.8) with gentle inversion. Samples were then cryoprotected in 20% sucrose, embedded in OCT, and sectioned at 30 µm. Sections were collected on poly-L-lysine–coated slides (Ted Pella, Prod. No. 18026) and stored at −80 °C until atomic force microscopy (AFM) measurements.

Prior to AFM analysis, sections were briefly thawed, washed in phosphate buffer, and kept hydrated in a humidity chamber using a hydrophobic barrier pen. AFM measurements were performed using a Bruker Dimension Icon microscope equipped with Bruker SPM-SPH 5µm spherical probes. Thermal noise calibration was performed before each measurement. For each sample, 3–12 nanoindentations were acquired at three distinct locations within the lesion core.

Raw deflection–height data were converted to force–indentation curves and fit in *Bruker NanoScope Analysis* software to the Hertz–Spherical contact model:

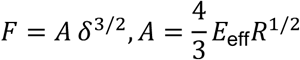

where *F* is the applied force, *δ* is the indentation depth, *Eₑff* is the effective Young’s modulus, and *R* is the probe radius. The apparent tissue Young’s modulus (*Eₛ*) was then derived from the fitting constant *A* according to:

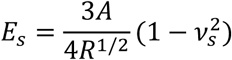

where *νₛ* is the Poisson’s ratio of the sample (0.4). Stiffness values (*Eₛ*) were reported in Pascals (Pa).

### Body curvature analysis

Adult wild type and *gata4^wcm6/wcm6^* fish were briefly anesthetized in 0.02% aminobenzoic-acid-ethyl methyl-ester (MS222, Sigma) and lateral images were obtained before SCI and subsequently at 2, 4 and 6 wpi. Curvature was calculated in MATLAB using the Curvature function of the *Curvature of a 1D curve in a 2D or 3D space* script^67^. Curvature analysis was performed blinded and the XY coordinates from the dorsal line between the operculum and the caudal fin were taken for each individual fish.

### Single-Nucleus RNA sequencing

Spinal cord segments (2.0 mm anterior and posterior to the injury site) were dissected from *gata4^wcm6/wcm6^* and wild-type fish at 2 wpi, flash-frozen, and stored at −80 °C. Nuclei isolation was adapted from a previously established protocol ⁵¹. All solutions, reagents, and tools were pre-chilled and maintained on ice throughout the procedure. Nuclei Isolation Buffer 1 (NIM1) consisted of 250 mM sucrose (Sigma, S0389), 25 mM KCl, 5 mM MgCl₂, and 10 mM Tris (pH 8.0; Fisher, 10376743) prepared in RNase-free water. Nuclei Isolation Buffer 2 (NIM2) was prepared by supplementing NIM1 with 1 µM DTT, 0.4 U/µL RNase inhibitor (New England Biolabs, M0307S), and 0.1% Triton X-100 (Sigma, CAS 77861). A swinging-bucket centrifuge was pre-cooled to 4 °C prior to sample processing. Tissue samples were incubated in cold NIM2 buffer and gently dissociated by trituration on ice, followed by centrifugation at 200 × g for 10 minutes at 4 °C. The resulting pellet was resuspended in Nuclei Buffer containing 0.75% bovine serum albumin and passed through a 40 µm cell strainer into a 15-mL conical tube coated with 1% bovine serum albumin. Following filtration, Hoechst dye (1 µg/mL) and Nuclei Fixation Solution were added, and samples were incubated on ice for 10 minutes. Nuclei were fluorescence-activated cell sorted (FACS) in 1× phosphate buffer (approximately 1 × 10⁶ nuclei per sample), transferred to a new 1% bovine serum albumin–coated 15-mL conical tube, and centrifuged at 200 × g for 10 minutes at 4 °C. The supernatant was carefully removed, leaving approximately 1.0 mL above the pellet. Nuclei were permeabilized by adding Nuclei Permeabilization Solution and incubating on ice for 3 minutes, followed by neutralization with Nuclei Neutralization Buffer and centrifugation at 200 × g for 10 minutes at 4 °C. The final pellet was fully resuspended in Nuclei Buffer supplemented with DMSO and stored at −80 °C until library preparation. Single-nucleus RNA-sequencing libraries were generated using the PARSE Evercode Whole Transcriptome Mini v2 kit according to the manufacturer’s instructions.

### Single-Nucleus RNA sequencing analysis

Raw-counts matrices were read into Seurat (v.5) and filtered to cells containing over 500 unique detected genes. One wild-type replicate had poor quality and was excluded from the analyses. Standard normalization, feature selection, and scaling procedures were employed. After principal component analysis and dimension reduction via uniform manifold approximation and projection, cells were annotated by cluster as follows: fibroblasts (*pdgfb*, *col1a1b*), ependymo-radial glia (*gfap*, *sox2*, *slc4a10a*), glutamatergic neurons (*snap25a*, *elavl3*), CSF–contacting neurons (*pkd2l1*), serotonergic neurons (*tph2*), oligodendrocyte precursor cells (*cspg4*, *sema5a*), oligodendrocytes (*mpz*, *mag*), microglia / macrophages (*ptprc*, *mpeg1.1*, *itage.1*, *mrc1b*), T cells (*lck*, *runx3*), and proliferating immune cells (*mki67*, *ptprc*). Within ERGs, differentially expressed genes were computed using a Wilcoxon test and Bonferroni post-hoc adjustment, as implemented in Seurat. Gene set enrichment, based on genes rank ordered by fold change, was computed using the WebGestalt online tool, using the KEGG pathways.

Mouse datasets for both spinal cord astrocytes^35^ and fibroblasts^42^ were obtained and opened in Seurat. The selected cell populations were then analyzed identically to the zebrafish ependymo-radial glial cells, namely, differential expression via Wilcoxon testing and gene set enrichment analysis via WebGestalt online tools, based on fold change rank ordering.

### Motif analyses

The position probability matrix for each TF was pulled from the JASPAR vertebrate library and compiled into a dataset which was then fed into FIMO. For this analysis we filtered FIMO hits selecting for p<0.001. This FIMO output was used as the primary dataset for motif analysis. To visualize TF binding site clusters, a custom python script (plot_fimo_clusters.py) was developed. This script reads the FIMO output, computes the midpoint of each hit, and converts it into a coordinate system relative to the *gata4* TSS. Because *gata4* is located on the minus strand, upstream regions are represented as positive distances from the TSS (0) along the x-axis. To identify local motif clusters, the script performs a sliding-window analysis across the upstream regulatory region using a 250 bp window with a 50 bp step size. For each window, the script calculates the total number of motif hits, the number of distinct TF families represented, and the lowest q-value among all hits within the window. Windows were classified as cluster-like if they contained more than four motif hits, represented more than three distinct TF families, and included at least one hit with a q-value < 0.05.

### Tissue bridge measurement

Tissue bridge diameter was quantified in ImageJ/FIJI. For each fish, 3–5 20 μm sections containing the thickest portion of the tissue bridge were analyzed. Lesion diameter was measured and normalized to the uninjured spinal cord diameter caudal to the lesion.

### Loxl2 immunostaining quantification

A 300 µm region of interest (ROI) was centered on the spinal cord stump apex, guided by GFAP signal. The FIJI “Analyze Particles” function was used to determine the percentage area occupied by Loxl2 within the ROI. Measurements were performed blinded, using at least three sections per fish, and the values were averaged to yield a single value for each animal.

### Confocal imaging and mean gray value analysis

Spinal cord images of TMR-O– and TMR-NB–treated fish were acquired under identical confocal settings (Galvano scan mode; integration = 1; dwell time = 0.4 µs; 561 nm laser at 0.639%; z-step = 3.4 µm; 7 z-steps). Maximum intensity projections were generated in FIJI. A polygonal ROI was drawn around the lesion site, excluding bone, and Mean Gray Value was measured using the FIJI “Measure” function. For each animal, 2–5 sections were analyzed.

### Co-localization analyses

For non-nuclear colocalization analyses, images from 2–5 sections per animal were processed in Nikon NIS-Elements AR. Within each measurement set, channels were uniformly thresholded to generate binary masks for each signal. Manders’ colocalization coefficients were then calculated from the thresholded images, multiplied by 100, and reported as percent overlap.

For nuclear-based colocalization measurements (e.g., EdU, PCNA, Sox2), nuclei within 450 µm of the lesion site were manually counted across 2–5 sections per animal in Nikon NIS-Elements AR (Edu^+^EGFP^+^) or Fiji (PCNA^+^Sox2^+^), and the proportion of nuclei overlapping the signal of interest was quantified.

### In Situ Hybridization quantification

In situ hybridization signal was quantified from 2–5 sections per animal using a custom Python script that extracted mean RGB values for each image within a defined ROI. Average intensity values for the red, green, and blue channels were calculated for each sample. RGB triplets were plotted in 3D color space to visualize distribution patterns. A consistent inverse correlation was observed between signal strength and red + green (R+G) intensity, reflecting the deeper blue/purple chromogenic product. Principal component analysis confirmed that R+G intensity captured the primary variance associated with in situ hybridization signal. Thus, R+G values were used for subsequent statistical comparisons.

### AFOG staining scoring

AFOG-stained sections (2–5 per animal) were imaged and scored blindly by two investigators and assigned to four bins: 0 = no detectable aniline blue staining, 1 = minimal staining, 2 = moderate staining, and 3 = maximal staining.

### Statistical tests

Spinal cord transection surgeries and regeneration analyses were performed in a blinded manner whenever possible. Experiments were repeated 2–4 times using independent clutches of animals. For violin plots, solid lines denote medians and dotted lines denote the 25th and 75th percentiles. All individual data points are shown in the graphs, with each point representing a single animal. Sample sizes, statistical tests, and P values are provided in the main figures or figure legends. All statistical tests were performed using GraphPad Prism. Where two groups are compared, data were assessed for normality, and either Welch’s *t*-test or the nonparametric Mann–Whitney test was used as appropriate. Where three or more groups were compared, one-way ANOVA with appropriate corrections for multiple comparisons were used.

## ACKNOWLEDGMENTS

We thank Ken Poss for supporting preliminary studies on Gata4 in his laboratory. We are especially grateful to Chris V.E. Wright for insightful and in-depth discussions and comments on the manuscript. Members of the Cigliola laboratory provided valuable feedback and discussion throughout the project and on the manuscript. We also thank Guillaume Dupuy and Cory Guthrie for zebrafish care, and Todd Evans for sharing the *gata4^wcm6/wcm6^* zebrafish line. We are grateful to Dimitry Koktysh at the Vanderbilt Institute of Nanoscale Science and Engineering (VINSE) for assistance with atomic force microscopy experiments, Agnès Loubat at the Flow Cytometry Platform of the Institute of Biology Valrose (Nice, France) for support with single-nucleus RNA-sequencing library preparation, and Spencer Dunn for assistance with computational tool development. This work was supported by research support from Vanderbilt University, the Vanderbilt Brain Institute, VINSE pilot funding, the French National Research Agency (ANR- 22-CPJ1 - 0045–01), the LABEX SIGNALIFE (ANR- 11-LABX- 0028–01) and the IDEX UCA Jedi program (ANR- 15-IDEX- 01) to V.C.

## REFERENCES

1. Bradbury, E.J., and Burnside, E.R. (2019). Moving beyond the glial scar for spinal cord repair. Nat Commun 10, 3879. 10.1038/s41467-019-11707-7.

2. Tran, A.P., Warren, P.M., and Silver, J. (2018). The Biology of Regeneration Failure and Success After Spinal Cord Injury. Physiol Rev 98, 881–917. 10.1152/physrev.00017.2017.

3. Soderblom, C., Luo, X., Blumenthal, E., Bray, E., Lyapichev, K., Ramos, J., Krishnan, V., Lai-Hsu, C., Park, K.K., Tsoulfas, P., and Lee, J.K. (2013). Perivascular fibroblasts form the fibrotic scar after contusive spinal cord injury. J Neurosci 33, 13882–13887. 10.1523/JNEUROSCI.2524-13.2013.

4. Goritz, C., Dias, D.O., Tomilin, N., Barbacid, M., Shupliakov, O., and Frisen, J. (2011). A pericyte origin of spinal cord scar tissue. Science 333, 238–242. 10.1126/science.1203165.

5. Dias, D.O., Kim, H., Holl, D., Werne Solnestam, B., Lundeberg, J., Carlen, M., Goritz, C., and Frisen, J. (2018). Reducing Pericyte-Derived Scarring Promotes Recovery after Spinal Cord Injury. Cell 173, 153–165 e122. 10.1016/j.cell.2018.02.004.

6. Wanner, I.B., Anderson, M.A., Song, B., Levine, J., Fernandez, A., Gray-Thompson, Z., Ao, Y., and Sofroniew, M.V. (2013). Glial scar borders are formed by newly proliferated, elongated astrocytes that interact to corral inflammatory and fibrotic cells via STAT3-dependent mechanisms after spinal cord injury. J Neurosci 33, 12870–12886. 10.1523/JNEUROSCI.2121-13.2013.

7. Sofroniew, M.V. (2009). Molecular dissection of reactive astrogliosis and glial scar formation. Trends Neurosci 32, 638–647. 10.1016/j.tins.2009.08.002.

8. Amlerova, Z., Chmelova, M., Anderova, M., and Vargova, L. (2024). Reactive gliosis in traumatic brain injury: a comprehensive review. Front Cell Neurosci 18, 1335849. 10.3389/fncel.2024.1335849.

9. Klapka, N., Hermanns, S., Straten, G., Masanneck, C., Duis, S., Hamers, F.P., Muller, D., Zuschratter, W., and Muller, H.W. (2005). Suppression of fibrous scarring in spinal cord injury of rat promotes long-distance regeneration of corticospinal tract axons, rescue of primary motoneurons in somatosensory cortex and significant functional recovery. Eur J Neurosci 22, 3047–3058. 10.1111/j.1460-9568.2005.04495.x.

10. Stichel, C.C., Hermanns, S., Luhmann, H.J., Lausberg, F., Niermann, H., D’Urso, D., Servos, G., Hartwig, H.G., and Muller, H.W. (1999). Inhibition of collagen IV deposition promotes regeneration of injured CNS axons. Eur J Neurosci 11, 632–646. 10.1046/j.1460-9568.1999.00466.x.

11. Becker, T., Wullimann, M.F., Becker, C.G., Bernhardt, R.R., and Schachner, M. (1997). Axonal regrowth after spinal cord transection in adult zebrafish. J Comp Neurol 377, 577–595. 10.1002/(sici)1096-9861(19970127)377:4<577::aid-cne8>3.0.co;2-#.

12. Reimer, M.M., Sorensen, I., Kuscha, V., Frank, R.E., Liu, C., Becker, C.G., and Becker, T. (2008). Motor neuron regeneration in adult zebrafish. J Neurosci 28, 8510–8516. 10.1523/JNEUROSCI.1189-08.2008.

13. Mokalled, M.H., Patra, C., Dickson, A.L., Endo, T., Stainier, D.Y., and Poss, K.D. (2016). Injury-induced ctgfa directs glial bridging and spinal cord regeneration in zebrafish. Science 354, 630–634. 10.1126/science.aaf2679.

14. Cigliola, V., Shoffner, A., Lee, N., Ou, J., Gonzalez, T.J., Hoque, J., Becker, C.J., Han, Y., Shen, G., Faw, T.D., et al. (2023). Spinal cord repair is modulated by the neurogenic factor Hb-egf under direction of a regeneration-associated enhancer. Nat Commun 14, 4857. 10.1038/s41467-023-40486-5.

15. Kolb, J., Tsata, V., John, N., Kim, K., Mockel, C., Rosso, G., Kurbel, V., Parmar, A., Sharma, G., Karandasheva, K., et al. (2023). Small leucine-rich proteoglycans inhibit CNS regeneration by modifying the structural and mechanical properties of the lesion environment. Nat Commun 14, 6814. 10.1038/s41467-023-42339-7.

16. Wehner, D., Tsarouchas, T.M., Michael, A., Haase, C., Weidinger, G., Reimer, M.M., Becker, T., and Becker, C.G. (2017). Wnt signaling controls pro-regenerative Collagen XII in functional spinal cord regeneration in zebrafish. Nat Commun 8, 126. 10.1038/s41467-017-00143-0.

17. Tsata, V., Mollmert, S., Schweitzer, C., Kolb, J., Mockel, C., Bohm, B., Rosso, G., Lange, C., Lesche, M., Hammer, J., et al. (2021). A switch in pdgfrb(+) cell-derived ECM composition prevents inhibitory scarring and promotes axon regeneration in the zebrafish spinal cord. Dev Cell 56, 509–524 e509. 10.1016/j.devcel.2020.12.009.

18. John, N., Fleming, T., Kolb, J., Lyraki, O., Vasquez-Sepulveda, S., Parmar, A., Kim, K., Tarczewska, M., Gupta, P., Singh, K., et al. (2025). Biphasic inflammation control by fibroblasts enables spinal cord regeneration in zebrafish. Cell Rep 44, 116469. 10.1016/j.celrep.2025.116469.

19. Goldshmit, Y., Sztal, T.E., Jusuf, P.R., Hall, T.E., Nguyen-Chi, M., and Currie, P.D. (2012). Fgf-dependent glial cell bridges facilitate spinal cord regeneration in zebrafish. J Neurosci 32, 7477–7492. 10.1523/JNEUROSCI.0758-12.2012.

20. Kikuchi, K., Holdway, J.E., Werdich, A.A., Anderson, R.M., Fang, Y., Egnaczyk, G.F., Evans, T., Macrae, C.A., Stainier, D.Y., and Poss, K.D. (2010). Primary contribution to zebrafish heart regeneration by gata4(+) cardiomyocytes. Nature 464, 601–605. 10.1038/nature08804.

21. Malek Mohammadi, M., Kattih, B., Grund, A., Froese, N., Korf-Klingebiel, M., Gigina, A., Schrameck, U., Rudat, C., Liang, Q., Kispert, A., et al. (2017). The transcription factor GATA4 promotes myocardial regeneration in neonatal mice. EMBO Mol Med 9, 265–279. 10.15252/emmm.201606602.

22. Yu, W., Huang, X., Tian, X., Zhang, H., He, L., Wang, Y., Nie, Y., Hu, S., Lin, Z., Zhou, B., et al. (2016). GATA4 regulates Fgf16 to promote heart repair after injury. Development 143, 936–949. 10.1242/dev.130971.

23. Arroyo, N., Villamayor, L., Diaz, I., Carmona, R., Ramos-Rodriguez, M., Munoz-Chapuli, R., Pasquali, L., Toscano, M.G., Martin, F., Cano, D.A., and Rojas, A. (2021). GATA4 induces liver fibrosis regression by deactivating hepatic stellate cells. JCI Insight 6. 10.1172/jci.insight.150059.

24. Delgado, I., Carrasco, M., Cano, E., Carmona, R., Garcia-Carbonero, R., Marin-Gomez, L.M., Soria, B., Martin, F., Cano, D.A., Munoz-Chapuli, R., and Rojas, A. (2014). GATA4 loss in the septum transversum mesenchyme promotes liver fibrosis in mice. Hepatology 59, 2358–2370. 10.1002/hep.27005.

25. Winkler, M., Staniczek, T., Kurschner, S.W., Schmid, C.D., Schonhaber, H., Cordero, J., Kessler, L., Mathes, A., Sticht, C., Nessling, M., et al. (2021). Endothelial GATA4 controls liver fibrosis and regeneration by preventing a pathogenic switch in angiocrine signaling. J Hepatol 74, 380–393. 10.1016/j.jhep.2020.08.033.

26. Zhang, Z., Shayani, G., Xu, Y., Kim, A., Hong, Y., Feng, H., and Zhu, H. (2023). Induction of Senescence by Loss of Gata4 in Cardiac Fibroblasts. Cells 12. 10.3390/cells12121652.

27. Agnihotri, S., Wolf, A., Picard, D., Hawkins, C., and Guha, A. (2009). GATA4 is a regulator of astrocyte cell proliferation and apoptosis in the human and murine central nervous system. Oncogene 28, 3033–3046. 10.1038/onc.2009.159.

28. Agnihotri, S., Wolf, A., Munoz, D.M., Smith, C.J., Gajadhar, A., Restrepo, A., Clarke, I.D., Fuller, G.N., Kesari, S., Dirks, P.B., et al. (2011). A GATA4-regulated tumor suppressor network represses formation of malignant human astrocytomas. J Exp Med 208, 689–702. 10.1084/jem.20102099.

29. Heicklen-Klein, A., and Evans, T. (2004). T-box binding sites are required for activity of a cardiac GATA-4 enhancer. Dev Biol 267, 490–504. 10.1016/j.ydbio.2003.09.042.

30. Zhou, L., McAdow, A.R., Yamada, H., Burris, B., Klatt Shaw, D., Oonk, K., Poss, K.D., and Mokalled, M.H. (2023). Progenitor-derived glia are required for spinal cord regeneration in zebrafish. Development 150. 10.1242/dev.201162.

31. Klatt Shaw, D., Saraswathy, V.M., Zhou, L., McAdow, A.R., Burris, B., Butka, E., Morris, S.A., Dietmann, S., and Mokalled, M.H. (2021). Localized EMT reprograms glial progenitors to promote spinal cord repair. Dev Cell 56, 613–626 e617. 10.1016/j.devcel.2021.01.017.

32. Sam, J., Mercer, E.J., Torregroza, I., Banks, K.M., and Evans, T. (2020). Specificity, redundancy and dosage thresholds among gata4/5/6 genes during zebrafish cardiogenesis. Biol Open 9. 10.1242/bio.053611.

33. Yang, J., Savvatis, K., Kang, J.S., Fan, P., Zhong, H., Schwartz, K., Barry, V., Mikels-Vigdal, A., Karpinski, S., Kornyeyev, D., et al. (2016). Targeting LOXL2 for cardiac interstitial fibrosis and heart failure treatment. Nat Commun 7, 13710. 10.1038/ncomms13710.

34. Yamauchi, M., and Sricholpech, M. (2012). Lysine post-translational modifications of collagen. Essays Biochem 52, 113–133. 10.1042/bse0520113.

35. Li, C., Wu, Z., Zhou, L., Shao, J., Hu, X., Xu, W., Ren, Y., Zhu, X., Ge, W., Zhang, K., et al. (2022). Temporal and spatial cellular and molecular pathological alterations with single-cell resolution in the adult spinal cord after injury. Signal Transduct Target Ther 7, 65. 10.1038/s41392-022-00885-4.

36. Cheng, F., Shen, Y., Mohanasundaram, P., Lindstrom, M., Ivaska, J., Ny, T., and Eriksson, J.E. (2016). Vimentin coordinates fibroblast proliferation and keratinocyte differentiation in wound healing via TGF-beta-Slug signaling. Proc Natl Acad Sci U S A 113, E4320–4327. 10.1073/pnas.1519197113.

37. Torres, S., Garcia-Palmero, I., Herrera, M., Bartolome, R.A., Pena, C., Fernandez-Acenero, M.J., Padilla, G., Pelaez-Garcia, A., Lopez-Lucendo, M., Rodriguez-Merlo, R., et al. (2015). LOXL2 Is Highly Expressed in Cancer-Associated Fibroblasts and Associates to Poor Colon Cancer Survival. Clin Cancer Res 21, 4892–4902. 10.1158/1078-0432.CCR-14-3096.

38. Aumiller, V., Strobel, B., Romeike, M., Schuler, M., Stierstorfer, B.E., and Kreuz, S. (2017). Comparative analysis of lysyl oxidase (like) family members in pulmonary fibrosis. Sci Rep 7, 149. 10.1038/s41598-017-00270-0.

39. Rudman-Melnick, V., Adam, M., Stowers, K., Potter, A., Ma, Q., Chokshi, S.M., Vanhoutte, D., Valiente-Alandi, I., Lindquist, D.M., Nieman, M.L., et al. (2024). Single-cell sequencing dissects the transcriptional identity of activated fibroblasts and identifies novel persistent distal tubular injury patterns in kidney fibrosis. Sci Rep 14, 439. 10.1038/s41598-023-50195-0.

40. Sertie, A.L., Sossi, V., Camargo, A.A., Zatz, M., Brahe, C., and Passos-Bueno, M.R. (2000). Collagen XVIII, containing an endogenous inhibitor of angiogenesis and tumor growth, plays a critical role in the maintenance of retinal structure and in neural tube closure (Knobloch syndrome). Hum Mol Genet 9, 2051–2058. 10.1093/hmg/9.13.2051.

41. Sole-Boldo, L., Raddatz, G., Schutz, S., Mallm, J.P., Rippe, K., Lonsdorf, A.S., Rodriguez-Paredes, M., and Lyko, F. (2020). Single-cell transcriptomes of the human skin reveal age-related loss of fibroblast priming. Commun Biol 3, 188. 10.1038/s42003-020-0922-4.

42. Xue, X., Wu, X., Fan, Y., Han, S., Zhang, H., Sun, Y., Yin, Y., Yin, M., Chen, B., Sun, Z., et al. (2024). Heterogeneous fibroblasts contribute to fibrotic scar formation after spinal cord injury in mice and monkeys. Nat Commun 15, 6321. 10.1038/s41467-024-50564-x.

43. Moon, H.J., Finney, J., Xu, L., Moore, D., Welch, D.R., and Mure, M. (2013). MCF-7 cells expressing nuclear associated lysyl oxidase-like 2 (LOXL2) exhibit an epithelial-to-mesenchymal transition (EMT) phenotype and are highly invasive in vitro. J Biol Chem 288, 30000–30008. 10.1074/jbc.C113.502310.

44. Eyre, D.R., Paz, M.A., and Gallop, P.M. (1984). Cross-linking in collagen and elastin. Annu Rev Biochem 53, 717–748. 10.1146/annurev.bi.53.070184.003441.

45. Akam-Baxter, E.A., Bergemann, D., Ridley, S.J., To, S., Andrea, B., Moon, B., Ma, H., Zhou, Y., Aguirre, A., Caravan, P., et al. (2024). Dynamics of collagen oxidation and cross linking in regenerating and irreversibly infarcted myocardium. Nat Commun 15, 4648. 10.1038/s41467-024-48604-7.

46. Akam-Baxter, E., Ridley, S.J., Moon, B.F., Caravan, P., and Sosnovik, D.E. (2025). Collagen Pyridinoline Cross-Links Are Absent in Reversible Myocardial Fibrosis. Circ Res 136, 315–317. 10.1161/CIRCRESAHA.124.325406.

47. Mollmert, S., Kharlamova, M.A., Hoche, T., Taubenberger, A.V., Abuhattum, S., Kuscha, V., Kurth, T., Brand, M., and Guck, J. (2020). Zebrafish Spinal Cord Repair Is Accompanied by Transient Tissue Stiffening. Biophys J 118, 448–463. 10.1016/j.bpj.2019.10.044.

48. Rodriguez, H.M., Vaysberg, M., Mikels, A., McCauley, S., Velayo, A.C., Garcia, C., and Smith, V. (2010). Modulation of lysyl oxidase-like 2 enzymatic activity by an allosteric antibody inhibitor. J Biol Chem 285, 20964–20974. 10.1074/jbc.M109.094136.

49. Alper, S.R., and Dorsky, R.I. (2022). Unique advantages of zebrafish larvae as a model for spinal cord regeneration. Front Mol Neurosci 15, 983336. 10.3389/fnmol.2022.983336.

50. Becker, C.J., Cigliola, V., Gillotay, P., Rich, A., De Simone, A., Han, Y., Di Talia, S., and Poss, K.D. (2023). In toto imaging of glial JNK signaling during larval zebrafish spinal cord regeneration. Development 150. 10.1242/dev.202076.

51. Li, Y., Han, W., Wu, Y., Zhou, K., Zheng, Z., Wang, H., Xie, L., Li, R., Xu, K., Liu, Y., et al. (2019). Stabilization of Hypoxia Inducible Factor-1alpha by Dimethyloxalylglycine Promotes Recovery from Acute Spinal Cord Injury by Inhibiting Neural Apoptosis and Enhancing Axon Regeneration. J Neurotrauma 36, 3394–3409. 10.1089/neu.2018.6364.

52. Gerri, C., Marin-Juez, R., Marass, M., Marks, A., Maischein, H.M., and Stainier, D.Y.R. (2017). Hif-1alpha regulates macrophage-endothelial interactions during blood vessel development in zebrafish. Nat Commun 8, 15492. 10.1038/ncomms15492.

53. Jain, S.K., and Subrahmanyam, D. (1978). On the mechanism of phenylhydrazine-induced hemolytic anemia. Biochem Biophys Res Commun 82, 1320–1324. 10.1016/0006-291x(78)90332-7.

54. Jopling, C., Sune, G., Faucherre, A., Fabregat, C., and Izpisua Belmonte, J.C. (2012). Hypoxia induces myocardial regeneration in zebrafish. Circulation 126, 3017–3027. 10.1161/CIRCULATIONAHA.112.107888.

55. Saputra, F., Kishida, M., and Hu, S.Y. (2024). Oxidative stress induced by hydrogen peroxide disrupts zebrafish visual development by altering apoptosis, antioxidant and estrogen related genes. Sci Rep 14, 14454. 10.1038/s41598-024-64933-5.

56. Grant, C.E., Bailey, T.L., and Noble, W.S. (2011). FIMO: scanning for occurrences of a given motif. Bioinformatics 27, 1017–1018. 10.1093/bioinformatics/btr064.

57. Chia, N.Y., Deng, N., Das, K., Huang, D., Hu, L., Zhu, Y., Lim, K.H., Lee, M.H., Wu, J., Sam, X.X., et al. (2015). Regulatory crosstalk between lineage-survival oncogenes KLF5, GATA4 and GATA6 cooperatively promotes gastric cancer development. Gut 64, 707–719. 10.1136/gutjnl-2013-306596.

58. Hu, X., Li, T., Zhang, C., Liu, Y., Xu, M., Wang, W., Jia, Z., Ma, K., Zhang, Y., and Zhou, C. (2011). GATA4 regulates ANF expression synergistically with Sp1 in a cardiac hypertrophy model. J Cell Mol Med 15, 1865–1877. 10.1111/j.1582-4934.2010.01182.x.

59. Morin, S., Charron, F., Robitaille, L., and Nemer, M. (2000). GATA-dependent recruitment of MEF2 proteins to target promoters. EMBO J 19, 2046–2055. 10.1093/emboj/19.9.2046.

60. Mathison, M., Singh, V.P., Sanagasetti, D., Yang, L., Pinnamaneni, J.P., Yang, J., and Rosengart, T.K. (2017). Cardiac reprogramming factor Gata4 reduces postinfarct cardiac fibrosis through direct repression of the profibrotic mediator snail. J Thorac Cardiovasc Surg 154, 1601–1610 e1603. 10.1016/j.jtcvs.2017.06.035.

61. Gilad, G.M., and Gilad, V.H. (2001). Beta-aminopropionitrile treatment can accelerate recovery of mice after spinal cord injury. Eur J Pharmacol 430, 69–72. 10.1016/s0014-2999(01)01354-1.

62. Park, H.C., Kim, C.H., Bae, Y.K., Yeo, S.Y., Kim, S.H., Hong, S.K., Shin, J., Yoo, K.W., Hibi, M., Hirano, T., et al. (2000). Analysis of upstream elements in the HuC promoter leads to the establishment of transgenic zebrafish with fluorescent neurons. Dev Biol 227, 279–293. 10.1006/dbio.2000.9898.

63. Bernardos, R.L., and Raymond, P.A. (2006). GFAP transgenic zebrafish. Gene Expr Patterns 6, 1007–1013. 10.1016/j.modgep.2006.04.006.

64. Zhao, L., Borikova, A.L., Ben-Yair, R., Guner-Ataman, B., MacRae, C.A., Lee, R.T., Burns, C.G., and Burns, C.E. (2014). Notch signaling regulates cardiomyocyte proliferation during zebrafish heart regeneration. Proc Natl Acad Sci U S A 111, 1403–1408. 10.1073/pnas.1311705111.

65. Shin, J., Park, H.C., Topczewska, J.M., Mawdsley, D.J., and Appel, B. (2003). Neural cell fate analysis in zebrafish using olig2 BAC transgenics. Methods Cell Sci 25, 7–14. 10.1023/B:MICS.0000006847.09037.3a.

66. Ying, Y., Wu, J and He, M (2021). In-situ hybridization for mouse brain sections. Bio-protocol.

67. Mjaavatten, A. (2025). Curvature of a 1D curve in a 2D or 3D space. (https://www.mathworks.com/matlabcentral/fileexchange/69452-curvature-of-a-1d-curve-in-a-2d-or-3d-space) MATLAB Central File Exchange.

